# An integrated multi-omic single cell atlas to redefine human B cell memory

**DOI:** 10.1101/801530

**Authors:** David R. Glass, Albert G. Tsai, John Paul Oliveria, Felix J. Hartmann, Samuel C. Kimmey, Ariel A. Calderon, Luciene Borges, Sean C. Bendall

**Affiliations:** Immunology Graduate Program, Stanford University, Stanford, CA, 94305, USA; Department of Pathology, Stanford University, Stanford, CA, 94305, USA; Department of Medicine, Division of Respirology, McMaster University, Hamilton, ON, L8S4K1, Canada; Department of Developmental Biology, Stanford, University, Stanford CA, 94305, USA

**Author notes:** Co-first author (David R. Glass, Albert G. Tsai).

## Abstract

To evaluate the impact of heterogeneous B cells in health and disease, comprehensive profiling is needed at a single cell resolution. We developed a highly-multiplexed screen to quantify the co-expression of 351 surface molecules on low numbers of primary cells. We identified dozens of differentially expressed molecules and aligned their variance with B cell isotype usage, metabolism, biosynthesis activity, and signaling response. Here, we propose a new classification scheme to segregate peripheral blood B cells into ten unique subsets, including a CD45RB+ CD27- early memory population and a CD19^hi^ CD11c+ memory population that is a potent responder to immune activation. Furthermore, we quantify the contributions of antibody isotype and cell surface phenotype to various cell processes and find that phenotype largely drives B cell function. Taken together, these findings provide an extensive profile of human B cell diversity that can serve as a resource for further immunological investigations.

## Introduction

B cells have a profound effect on human health. Their ability to create a diverse antibody repertoire protects against an ever-evolving landscape of pathogens, while the formation of immunological memory suppresses repeat infection. A robust antibody response is the primary goal of nearly all current vaccinations and remains as the best correlate of vaccine efficacy (Plotkin, 2010). B cells have, however, also been implicated in pathogenic roles in autoimmunity (Magro, Clair and Tedder, 2008), transplant (Karahan, Claas and Heidt, 2017), and cancer (Sarvaria, Madrigal and Saudemont, 2017). Indeed, targeting B cells with anti-CD20 therapy in each of these diseases has shown promising results (Huynh et al. 2017; Sood and Hariharan 2018; Thaunat, Morelon, and Defrance 2010), though it is not always clear whether the therapeutic benefits arise from mitigation of antibody generation or disruption of some alternative B cell function.

Canonical gating strategies segregate B cells into five populations: transitional, naïve, non-switched memory, switched memory, and plasma cells (Maecker, 2012). Transitional cells (CD24^hi^ CD38^hi^ CD27- IgMD+) are recent bone marrow emigrants transitioning from immature to mature B cells. Naïve B cells (CD38^lo^/- CD27- IgMD+) are mature B cells that are ‘naïve’ to antigen and have never been stimulated by their cognate antigen. Memory cells (CD38^lo^/- CD27+) have taken part in previous immune responses and form part of the immunological memory of that event to prevent reinfection. These cells are further segregated as non-switched (IgM+ and/or IgD+) or switched IgM-, IgD-), based on their immunoglobulin heavy chain (IgH) isotypes which can ‘switch’ from the immature isotypes (IgM and IgD) to the mature isotypes (IgG, IgA, and IgE) after stimulation. Each antibody isotype confers unique downstream effector functions to membrane-bound and secreted antibodies and isotype usage can have both tissue- and pathogen-specific associations (Janeway *et al*., 2005). Plasma cells (CD38^hi^ CD27^hi^) also arise after immune activation as effector cells that secrete large quantities of antibodies.

This canonical gating strategy for B cells lacks the granularity and functional segregation that is captured in canonical T cell subsetting (Maecker, 2012). Dozens of T cell subsets can be gated using cell surface phenotypic identifiers that serve as proxies for cytotoxicity, cytokine production, activation status, and immunoregulatory capabilities. Furthermore, discrimination of naïve and memory B cells is based on expression of CD27, but a number of studies have reported CD27- memory phenotypes (Thorarinsdottir *et al*., 2016; Karnell *et al*., 2017). IgH isotype is often used to augment B cell gating (Berkowska *et al*., 2011; Krishnamurty *et al*., 2016), but others have suggested that functional differences are better captured with phenotypic subsetting rather than isotypic subsetting (Zuccarino-Catania *et al*., 2014). These discrepancies highlight our inability to consistently identify and sort functional subsets of human B cells, impeding our capacity to selectively target pathogenic B cells in autoimmunity and induce memory responses in vaccination.

To identify optimal features for B cell subsetting and prospective isolation, many parameters must be quantified across a large number of B cells to ensure that even rare populations are accounted for. This is particularly important in the context of disease, as cell frequencies can alter dramatically in immune activation and/or dysregulation (Hartmann *et al*., 2019), so populations that are rare in homeostasis may expand in disease. The application of next-generation sequencing (NGS) to basic B cell biology has revealed insights into isotype class-switching (Horns *et al*., 2016), self/foreign recognition (Burnett *et al*., 2018), and antibody evolution (Croote *et al*., 2018), but functional and phenotypic B cell profiles to parallel this remain elusive. Functional activity is primarily the result of protein-protein interactions, and as mRNA abundance is an inconsistent predictor of protein abundance (Peterson *et al*., 2017), a proteomic-based approach is desirable to identify functional subsets of B cells. Mass cytometry (CyTOF) facilitates routine quantification of 40+ proteins on millions of single cells in a single experiment (Bendall et al. 2011) and has also been employed for functional readouts such as post-translational modifications (Levine *et al*., 2015), cell division (Good *et al*., 2019), cytokine production (Hartmann *et al*., 2016), and immune antigen-specificity (Newell *et al*., 2012).

We previously applied mass cytometry to describe the initial stages of B cell development that occur in the continuous differentiation of hematopoietic stems cells into immature B cells (Bendall *et al*., 2014). Here, to capture the further diversification of phenotype and function that occur in mature B cells in the periphery, we developed a highly-multiplexed single cell screen to quantify the co-expression of 351 surface molecules using mass cytometry. We identified 98 molecules expressed on peripheral B cells, several of which can be used to segregate unique B cell subsets, including a CD45RB+ CD27- early memory population and a CD19^hi^ CD11c+ memory population that is a potent responder to immune activation. Here, we propose a new classification scheme that subsets peripheral B cells into ten unique populations and provide extensive single cell profiles of cell surface phenotype, isotype usage, metabolism, biosynthesis activity, and signaling response to immune activation. Furthermore, we assess the relative contribution of antibody isotype and cell phenotype to various cell processes and find that phenotype largely drives B cell function. This atlas of human B cell diversity will enable further studies to interrogate functional B cell subsets in the context of homeostasis, vaccination, infection, autoimmunity, and cancer.

## Results

### A highly-multiplexed single cell surface screen reveals the human B cell surface proteome

To identify molecules that could potentially differentiate unique B cell subsets, we developed a highly-multiplexed cell surface screen and quantified the co-expression of 351 surface antigens on healthy, human B cells (n=2, Figure 1A). We designed 12 mass cytometry antibody panels, each consisting of nine conserved molecules (CD45, CD7, CD11b, CD19, CD24, CD38, CD27, IgM, and IgD) for subsetting, and ∼30 variable molecules that were unique to each panel, for discovery (Table S1). The conserved molecules were selected such that cells analyzed with each of the 12 panels could be gated into four canonical B cell subsets found in peripheral blood: transitional, naïve, non-switched memory, and switched memory (Figure S1A, Figure 1B). Plasma cells were left out of the analysis due to insufficient numbers in healthy subjects after cryopreservation. The expression of the variable molecules across all panels was assessed for each canonical population. The conserved molecules were also used to generate a low-dimensional representation of the high-dimensional data using UMAP (Becht et al., 2019), facilitating visualization of co-expression patterns of variable molecules in an unbiased manner (see *Methods*).

**Figure 1:**
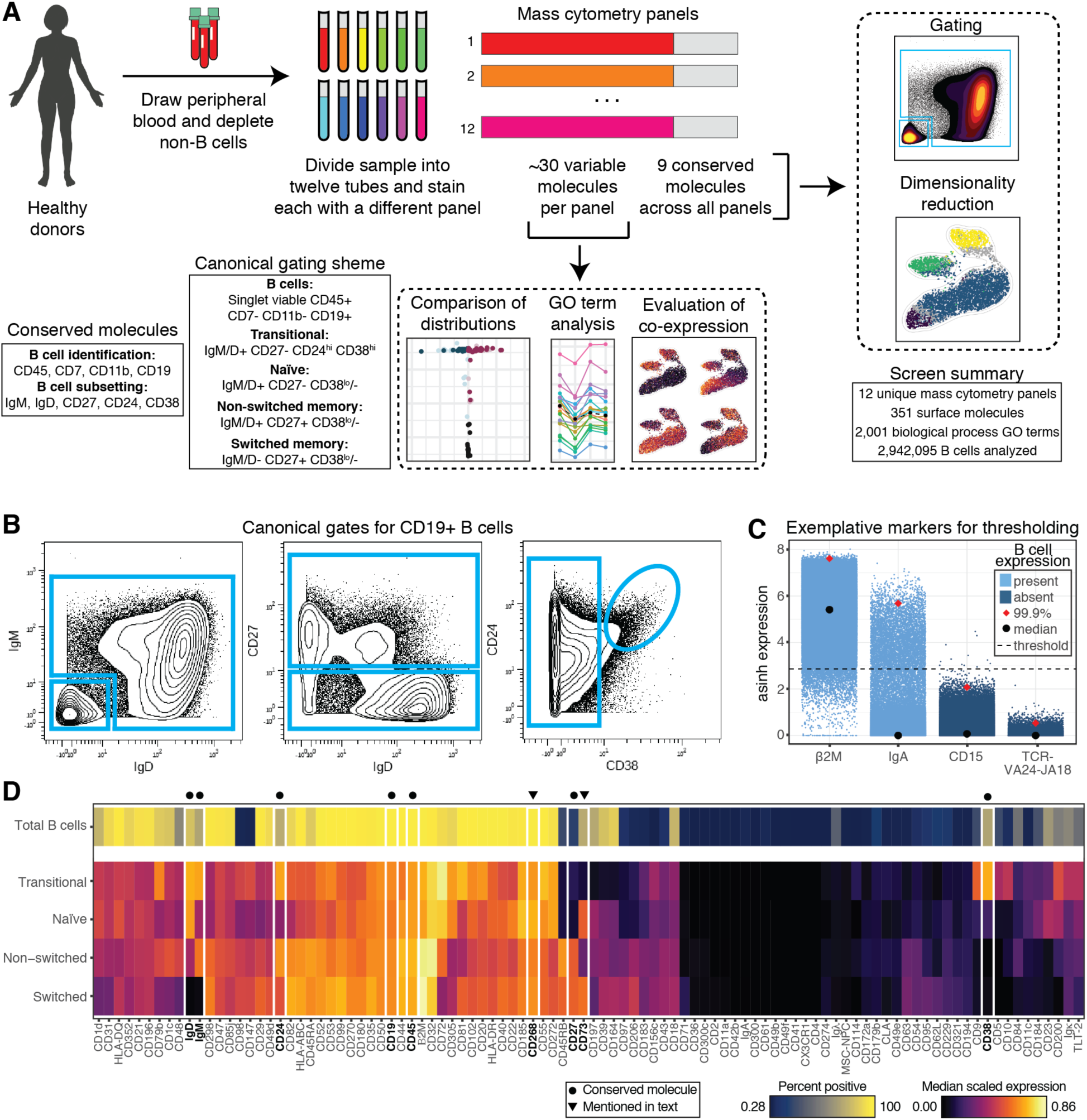
A highly-multiplexed single cell surface screen reveals the human B cell surface proteome. **A)** Experimental workflow for and summary of surface screen. Peripheral blood was drawn from healthy, human donors (n=2), density gradient centrifugation was performed, and cells were aliquoted and frozen. Aliquots were thawed, magnetically depleted of non-B cells, and stained with one of twelve custom-designed mass cytometry panels, each consisting of ∼40 metal-tagged antibodies. Each panel contained nine conserved molecules used for manual gating of canonical B cell subsets and dimensionality reduction. The remaining ∼30 molecules on each panel were screened for their co-expression on B cells. **B)** Representative mass cytometry plots of the gating strategy for canonical B cell populations. Each plot includes total B cells, gated as singlet viable CD45+ CD7- CD11b- CD19+. **C)** Single cell asinh-transformed expression values of four exemplative molecules in donor-pooled total B cells. Black dot indicates median expression; red diamond indicates 99.9^th^ percentile expression; dotted line indicates threshold of positivity (asinh-transformed value of 2.78). Molecules in light blue were considered present on B cells and used for downstream analysis. Molecules in dark blue were considered absent on B cells and removed from downstream analysis. **D)** For each molecule determined to be positive on B cells, percent positive of total B cells (top row) and median scaled expression of canonical B cell subsets (bottom four rows) are shown. The x-axis is hierarchically clustered based on median expression values. The y-axis is arranged in order of maturation (top to bottom). Molecules conserved in all mass cytometry panels in the screen are boxed, bolded, and noted with an dot. Molecules highlighted in the text are boxed, bolded, and noted with an arrow.

The collection of targets in the screen primarily consisted of surface molecules with immunology-associated gene ontology (GO) terms ontology (Boyle *et al*., 2004), including hundreds of CD molecules (Figure S1B). After setting a stringent threshold for classifying a molecules as present or absent on total, donor-pooled B cells using the 99.9^th^ percentile expression (see *Methods*, Figure 1C), we identified 98 surface molecules expressed on human B cells (Figure 1D). For each molecule, we quantified the percent positive in total B cells and the median scaled expression value in the four canonical subsets. Expression patterns between subsets recapitulated known biology (e.g. lower expression of IgD in non-switched memory cells compared to naïve cells; near uniform expression of the BAFF receptor, CD268), but also provided new insights. For example, the ecto-nucleotidase, CD73, was enriched in naïve and switched memory cells, while low/absent in transitional and non-switched memory cells. Interestingly, this molecule has been used to subset murine memory B cells (Tomayko *et al*., 2010), though its expression was mostly associated with *non-switched* memory cells in contrast to its expression in *switched* memory cells in our human dataset. CD73 is not typically used for B cell classification in humans, though it has been identified as a putative immunotherapy target in cancer due to its immunosuppressive functionality (Antonioli *et al*., 2016). Altogether, our single cell screen facilitated robust identification of surface molecules expressed by human B cells.

### Differential expression analysis reveals the anergic profile of naïve B cells

The canonical B cell gating scheme (Figure 1B), facilitated by the conserved molecules, organizes B cells by their state in a maturation process that occurs in the periphery (transitional --> naïve --> non-switched memory --> switched memory). To interrogate the changes in protein expression that occur throughout this process, we performed a Kolmogorov–Smirnov (KS) test between each pairwise combination of donor-pooled subsets, for all molecules (see *Methods*). We plotted the difference in median expression for the 61 significantly differentially-expressed molecules (p<0.005 after multiple hypothesis correction, Bonferroni method) and organized the heatmap such that molecules enriched in immature cells appear at the top and those enriched in mature cells appear at the bottom (Figure 2A).

**Figure 2:**
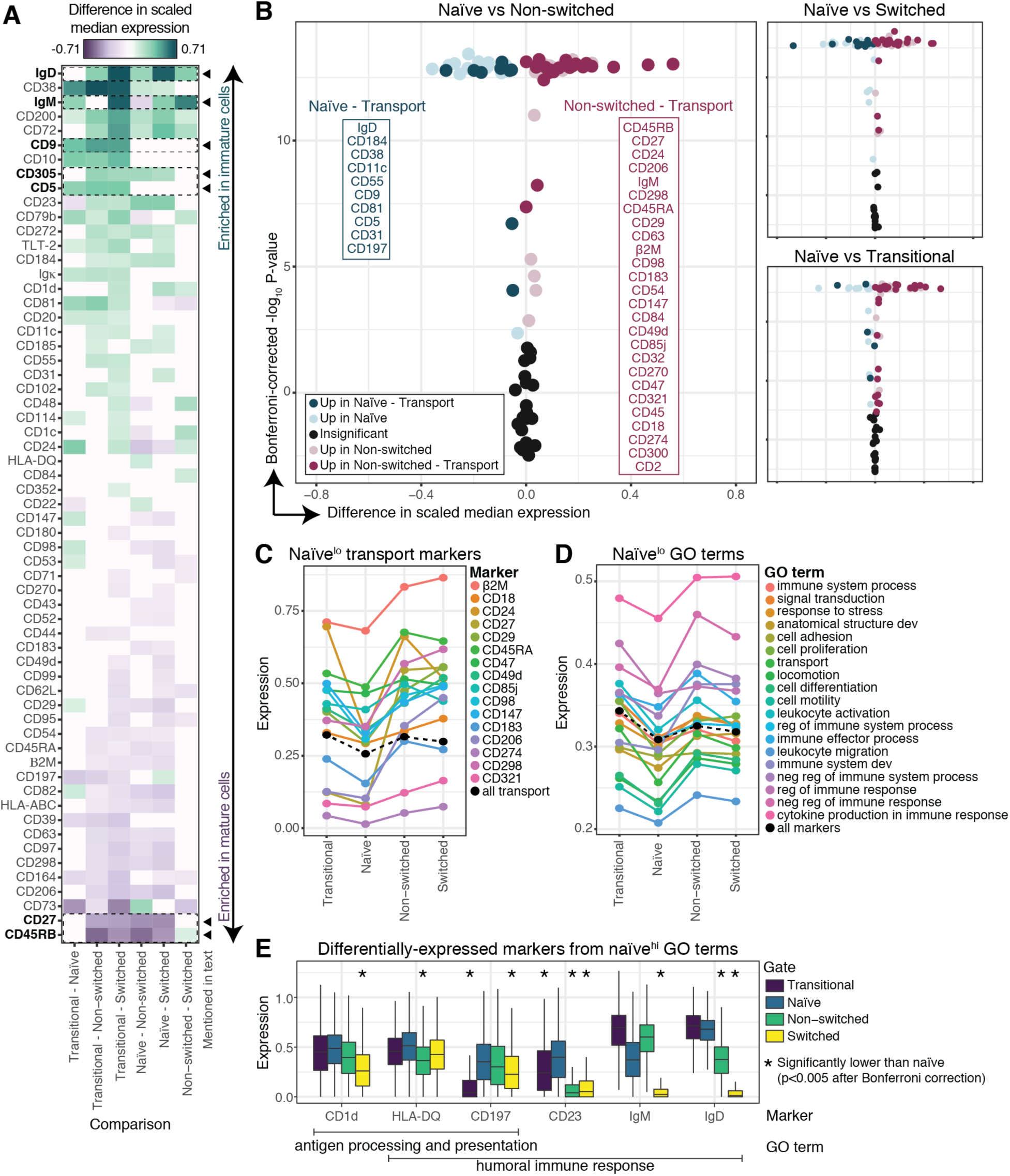
Differential expression analysis reveals the anergic profile of naïve B cells. **A)** Difference in scaled median expression for each pairwise comparison of canonical B cell subsets from pooled donors. Each column is a comparison of two B cell subsets, each row is a unique molecule. Only comparisons with an absolute difference > 0.1 and a p-value < 0.005 by KS test after Bonferroni correction are plotted – all other comparisons are absent from the heatmap or colored white. Rows are ordered by row-mean, resulting in an organization of the heatmap in which molecules enriched in immature cells appear at the top and those enriched in mature cells appear at the bottom. Molecules highlighted in the text are boxed, bolded, and indicated with an arrow. **B)** Volcano plot of the results of the comparisons visualized in A), colored by association with the GO term, transport. Molecules listed in boxes are those associated with transport and upregulated in the indicated subset, ordered by magnitude of difference of expression, from greatest to least (top to bottom). **C)** Median expression values of molecules associated with transport in which naïve cells have the lowest value (colors). Mean of the median expression values of all transport molecules (black). **D)** Mean of all median expression values of molecules associated with the indicated GO terms (color). Mean of all median expression values of all molecules expressed on B cells (black). **E)** Boxplots of expression values of six molecules significantly upregulated in naïve cells, and associated with “antigen processing and presentation” and/or “humoral immune response”. Naïve cells did not have the highest mean expression of any GO term amongst B cell subsets, but had the 2^nd^ highest mean expression for these two terms. Lines indicate median value, boxes indicate interquartile range (IQR), whiskers indicate first/third quartile +/- 1.5 IQR. Asterisks indicate populations with significantly lower expression than naïve cells (p < 0.005) by KS test with Bonferroni correction.

As expected, the immature isotypes, IgD and IgM, were enriched in antigen-inexperienced cells (transitional and naïve), while the canonical memory molecule, CD27 was enriched in memory cells. CD9, which has been reported to distinguish murine marginal zone, B-1, plasma (Won and Kearney, 2002), and/or regulatory (Sun *et al*., 2015) B cells was enriched in transitional cells over all other subsets; CD5, which has also been reported to distinguish murine B-1 (Berland and Wortis, 2002) and regulatory (Yanaba *et al*., 2008) B cells, showed similar patterning. CD305 (LAIR-1), which inhibits BCR signaling (Vries, 1999), was enriched in antigen-inexperienced cells over memory cells potentially increasing the antigen-specific activation threshold for these subsets. CD45RB, an isoform of CD45, displayed the opposite patterning and was more highly expressed in memory than in antigen-inexperienced cells. As with CD45RA/RO in naïve and memory T cells, CD45 isoforms are commonly definitional for functionally-distinct immune cell populations (Maecker, 2012) and have been reported as useful for B cell subsetting as well (Jackson *et al*., 2009).

We then asked if broader patterns in protein expression emerged from the pairwise comparison of B cell subsets. As an example, we plotted the results of our KS tests comparing naïve cells to the other three subsets and colored each molecule associated with the term “transport” (Figure 2B). The movement of substances or cellular components in and out of cells facilitates sending and receiving immune signals and is vital for coordinating the B cell response to an immune challenge. We found naïve cells expressed lower levels of molecules associated with transport than any other subset, suggesting they are less responsive to stimuli. Indeed, naïve cells had the lowest median expression value for 16 transport molecules and the lowest average expression across all 46 transport molecules (Figure 2C). We asked if this trend was consistent across GO terms and found that naïve cells had the lowest mean expression value for 19 out of 30 terms (Figure 2D). In fact, when the median expression value of all 98 molecules were averaged, naïve cells had the lowest expression value, suggesting they exist in a more anergic state than all other B cell subsets.

Since naïve cells downregulated most molecules on the screen relative to other subsets, we then asked if naïve cells were enriched for any GO terms. Naïve cells did not have the highest mean expression for any GO term and had the second highest mean expression for only two terms: “antigen processing and presentation” and “humoral immune response”. Within those two terms, only six molecules were significantly upregulated in naïve cells over at least one other subset – CD1d, HLA-DQ, CD197, CD23, IgM, and IgD (Figure 2E). In fact, in the entire screen, only CD23, the non-classical Fc receptor for IgE and IgG, was significantly upregulated in naïve cells over all other subsets. CD23 has been demonstrated to increase the threshold for B cell activation upon binding its ligands through induction of CD32, an inhibitory Fc receptor (Wang *et al*., 2015). These findings reveal there is a global downregulation of surface molecules in naïve cells coordinated with a specific upregulation of immunosuppressive molecules.

### CD45RB marks human memory B cells and identifies a unique early memory population

While CD27 is an accepted indication of B cell memory, even its earliest description indicated that there was already IgH rearrangement in the non-switched, CD27- pool (Klein, Rajewsky and Küppers, 1998). As such, to find markers that uniquely identify distinct B cells that are not captured by the canonical organizational scheme (Figure 1B), we analyzed the co-expression patterns of molecules in the screen across all B cells in an unbiased fashion. Because the surface screen was split across twelve reaction tubes, we could not directly determine whether a molecule expressed on a cell in tube X was co-expressed with a molecule in tube Y. We therefore generated a UMAP plot (Becht et al., 2019) organized by the expression of the conserved molecules (Figure 1A), using donor-pooled data from all twelve tubes (see *Methods*, Figure 3A). This two-dimensional representation of the high-dimensional data allowed us to visualize co-expression patterns of cells from different tubes in the same set of plots by overlaying molecule expression on UMAP coordinates.

**Figure 3:**
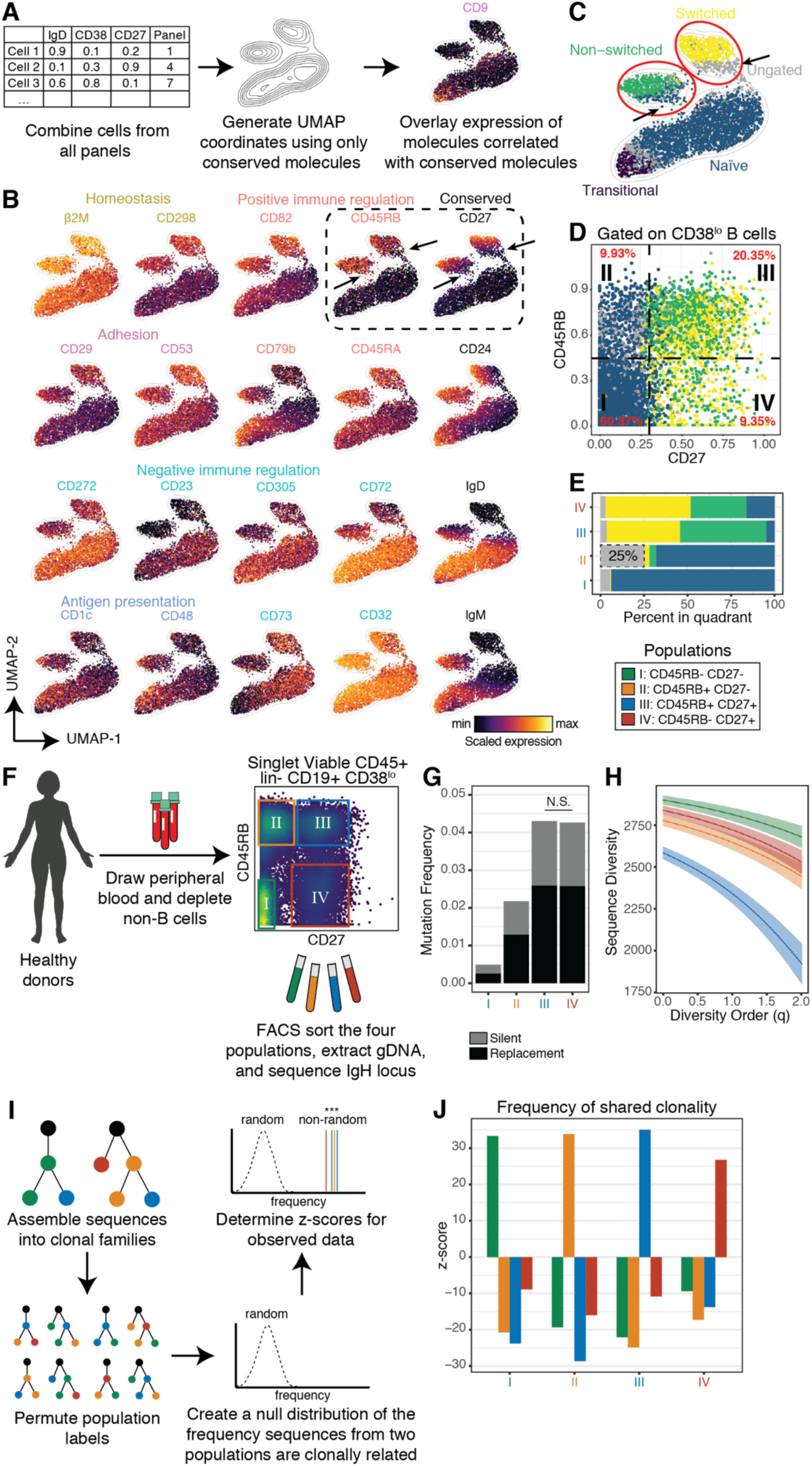
CD45RB marks human memory B cells and identifies a unique early memory population. **A)** Computational workflow for UMAP generation. UMAP coordinates were generated using conserved molecules as input on a random subset of donor-pooled cells from each of the twelve mass cytometry panels of the screen. **B)** UMAP plots colored by individually-scaled expression of the indicated molecule. Molecules are organized and colored by function or by their identity as a conserved molecule (right column). Molecules in each row are highly correlated (Pearson method, ± r > 0.3) with the conserved molecule in their row. The canonical memory marker (CD27) and putative memory marker (CD45RB) are boxed. Arrows indicate CD45RB+ CD27- cells. **C)** UMAP plot colored by canonical B cell subset. Circles indicate islands of cells with memory-like phenotypes based on UMAP co-localization and high-dimensional expression profiles. Arrows are the same as in B) and indicate cells with memory-like phenotypes that are not identified as memory cells by the canonical gating scheme. **D)** CD38^lo^ B cells (non-transitional B cells) are plotted on a biaxial and colored by their canonical B cell gate. Percent of CD38^lo^ B cells in each quadrant is quantified (red text). **E)** Percent of cells in each quadrant from D) by canonical B cell subset. High percentage of ungated cells in quadrant II is boxed and bolded. **F)** Experimental workflow for immune repertoire analysis. Peripheral blood was drawn from healthy, human donors (n=2), density gradient centrifugation was performed, and cell preparations were magnetically depleted of non-B cells. Singlet Viable CD45+ lin- CD19+ CD38^lo^ B cells were FACS-sorted from the four quadrants of CD27 x CD45RB, gDNA was extracted, and the IgH locus was sequenced by NGS. **G)** Mutation frequency of the IgH loci outside of the CDR3 region for sorted populations, colored by mutation type. Data are pooled from donors. All pairwise comparisons of populations were significantly different (Wilcoxon rank sum test with Bonferroni correction, p < 0.005), except where indicated. **H)** Sequence diversity (y-axis) of each sorted population across a range of diversity orders (x-axis). Shaded regions indicate 95% confidence interval, generated by bootstrapping. **I)** Computational workflow for clonal lineage analysis. Sequences were assembled into clonal lineages. Population labels for all sequences were randomly permuted and then for each population, the frequency in which a sequence from population X shared a clonal lineage with a sequence from population Y was quantified. This process was repeated 200 times to create a null distribution and then z-scores were derived for the observed data using the original population labels. **J)** Z-scores of observed data over a null distribution, relating the frequency a sequence from population X (x-axis) shared a clonal lineage with a sequence from population Y (colored bars).

Since expression of the conserved molecules undergo coordinated changes throughout B cell maturation, we hypothesized that other molecules correlated with the conserved molecules may also be useful for differentiating stages of B cell maturation. We plotted molecules that were highly correlated with the conserved molecules (Pearson method, ± r > 0.3) and organized by them function (label color) and correlated conserved marker (row, Figure 3B). The highest correlation (r=0.69) was between IgD and CD72, a negative regulator of B cell activation (Tsubata, 2012). IgD, which is highly-expressed on naïve cells, was also correlated with the negative regulators, CD23, CD305, and CD272 (Vendel *et al*., 2009). While IgM was correlated with CD32, the inhibitory Fc receptor, it was inversely correlated with the immunoregulatory molecule CD73, which we found was enriched in naïve and switched memory cells. CD27 was correlated with several molecules that positively regulate immune activation, potentially lowering the activation threshold of CD27+ cells.

We then asked how the canonical B cell fractions (Figure 1B - transitional, naïve, non-switched memory, switched memory, and ungated) related to the UMAP coordinates and overlaid expression patterns. Overlaying gating labels on the UMAP coordinates revealed that two “islands” in the plot that were not homogenously colored, indicating that while phenotypically similarly, these cells were considered different subsets by canonical gating (Figure 3C). The island with switched memory cells also contained ungated cells while the island with non-switched memory cells also contained both naïve and ungated cells (Figure 3C – arrows). Overlaying CD27 revealed that non-uniform expression of CD27 in these islands resulted in these mixed classifications (Figure 3B – arrows). Overlaying CD45RB, however, resulted in a more homogenous coloring of the two memory populations, while retaining an absence of expression in the transitional/naïve island. This suggested that CD45RB may have utility as a memory molecule. We plotted CD38lo/- cells (to remove transitional cells) on a CD27 x CD45RB biaxial and found that the majority of CD27+ cells were also CD45RB+ (RB+) and that the RB+ CD27- population contained 25% ungated cells (Figure 3D-E). Given the co-localization of RB+ CD27- cells and CD27+ cells on the UMAP, we hypothesized that that these cells represent a new population of memory cells, neglected under the current classification scheme.

To assess the spectrum of memory cell specification across the CD45RB and CD27 compartments, we prospectively isolated the four quadrants of the CD27 x CD45RB biaxial from healthy, human CD38lo/- B cells (n=2), extracted genomic DNA (gDNA), and sequenced the IgH loci by NGS for immune repertoire analysis (see *Methods*, Figure 3F). As a proxy for antigen exposure, we measured the donor-pooled mutation frequency of nucleotides in the IgH loci outside of the complementarity-determining region 3 (CDR3; Figure 3G; Boyd and Crowe, 2016). As expected, CD27+ cells had a relatively high mutational burden, acquired through somatic hypermutation (SHM) after exposure to antigen. Conversely, RB-CD27- cells showed a significantly lower mutational burden as they are still naïve to antigen (Wilcoxon rank sum test with Bonferroni correction, p<0.005). Interestingly, RB+ CD27- cells displayed an intermediate mutational burden, significantly higher than RB-CD27- cells and significantly lower than CD27+ cells. This level of mutational burden would be expected of an early population of memory cells, that have been exposed to antigen, but undergone fewer cycles of SHM than other memory cells.

After an immune challenge, B cells that are reactive to relevant antigens are selected to proliferate and differentiate into effector and memory classes. Naïve cells, therefore, tend to have more diverse immune repertoires as they have not undergone selection, while memory populations have less diverse immune repertoires (Briney *et al*., 2012). We quantified the diversity of the four populations across a range of diversity orders (Hafler *et al*., 2014) and found that RB-CD27- cells had the highest diversity, while RB+ CD27+ cells had the lowest diversity (see *Methods*, Figure 3H). Interestingly, both RB+ CD27- cells and RB-CD27+ cells had intermediate levels of diversity. This may indicate that RB+ CD27- cells undergo less stringent selection than RB+ CD27+ cells. This may also be true for RB-CD27+ cells, but given the high mutational burden of this population, it is also possible that this diversity is instead indicative of a more long-lived memory population that mediates protection against a lifetime of past immune challenges.

If expression of CD45RB was random and irrelevant to B cell activation and maturation, we would expect RB+ and RB-cells to share clonal lineages, as the molecule would not meaningfully segregate cells. We therefore asked if cells from one population tend to be clonally related to cells from any other population (see *Methods*, Figure 3I). We found that cells from each of the four populations were much more likely to share clonal lineages with cells from the same population than with those from a different population (Figure 3J). This indicates that expression of CD45RB and CD27 are highly coordinated within clonal lineages, as would be expected of two molecules that are upregulated in response to antigen engagement. Taken together, these findings provide strong evidence that expression of CD45RB is indicative of a peripheral blood memory B cell and, in conjunction with an absence of CD27, can be used to classify an early memory population.

### Segregating B cells into phenotypically and isotypically distinct subsets

The surface screen revealed several molecules differentially expressed in B cells and resulted in the identification of an early memory population (Figure 1-3), so we asked if we could better classify B cells into subsets by staining fresh, healthy, human peripheral blood B cells (n=3) with a mass cytometry panel comprised of the most informative B cell molecules from the screen (Figure 4A). To this end, we included canonical B cell molecules, heavy and light chain isotypes, and identified molecules with non-uniform expression on B cells (Table S1). To identify unique subsets, data was normalized and pooled from all donors (Figure S2A), and cells were first over-clustered with FlowSOM (Van Gassen *et al*., 2015) using all 40 molecules, then iteratively meta-clustered into ten distinct subsets (see *Methods*). IgH isotype was used for the initial over-clustering step, but not for subsequent meta-clustering to avoid creating artificial separations between phenotypically similar cells based on differential isotype usage alone.

**Figure 4:**
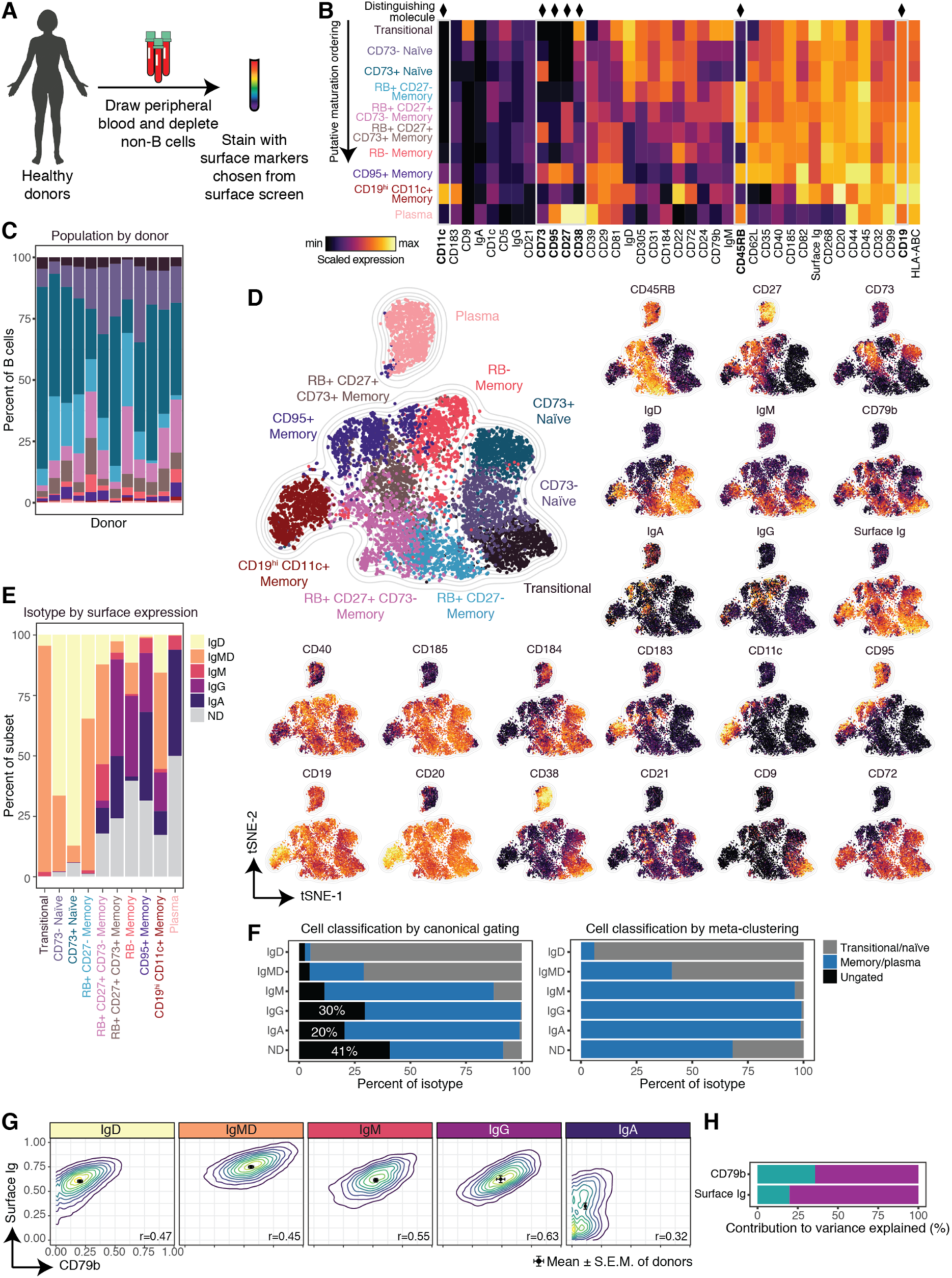
Segregating B cells into phenotypically and isotypically distinct subsets. **A)** Experimental workflow for phenotype discovery. Peripheral blood was drawn from healthy, human donors (n=3), density gradient centrifugation was performed, and cell preparations were magnetically depleted of non-B cells. Cells were stained with a mass cytometry panel consisting of informative molecules identified in the surface screen. **B)** Scaled median expression level for each molecule in each subset from pooled donors (range 0.00:0.91). X-axis is hierarchically clustered and y-axis is ordered in a putative arrangement of maturation (except for bottom three rows). Key molecules for distinguishing subsets are boxed, bolded, and noted with a diamond. **C)** Percent of B cells for all subsets segregated by donor from datasets analyzed in Figure 4 (n=3) and Figure 5 (n=9). Subsets are colored as in B). **D)** t-SNE plot generated from an equal subsampling of 1000 cells from each subset using only phenotypic (not isotypic) molecules. Cells are colored by subset identity or individually-scaled molecule expression. **E)** Percent of each B cell subset with a given IgH isotype class label (defined by surface staining of antibody isotypes). ND signifies “not determined” – cells with low/absent expression of all isotypes. IgMD denotes co-expression of IgM and IgD. **F)** Total B cells were segregated by heavy chain isotype. For each isotype, the percent of cells labeled as transitional/naïve, memory/plasma, and ungated was determined for the canonical gating scheme (left) and the meta-clustering classification (right). High percentages of ungated cells are bolded and enumerated. **G)** Contour plots of donor-pooled total B cells segregated by IgH isotype. Dots and error bar indicate mean and SEM of individual donors. Pearson correlations of CD79b and surface Ig expression levels are reported for each IgH isotype. **H)** Two linear models were created to predict single-cell expression of CD79b and surface Ig, respectively, using IgH isotype class labels and meta-clustering population labels as predictors. The relative contribution of phenotype (cyan) and isotype (magenta) to the variance explained by each model was quantified.

We then plotted the resulting surface expression profiles and arranged them in a putative order from least to most mature based on phenotypic similarity, IgH isotype usage, and expression of maturation molecules (Figure 4B, top to bottom). The characteristic expression pattern of seven distinguishing molecules were sufficient to manually gate each population and thus were also used to label the populations in this new scheme: CD11c, CD73, CD95, CD27, CD38, CD45RB, and CD19 (Figure S2B).

Two populations largely overlapped with canonical definitions: transitional (CD24^hi^ CD38^hi^ CD27-) and plasma (CD27^hi^ CD38^hi^) cells. Naïve cells (RB-CD27- CD38+/-) were further segregated into CD73+ and CD73-subsets. We identified six unique memory clusters, only two of which were uniformly CD27+. Two memory subsets were largely defined by expression of CD45RB and CD27: RB+ 27- (early) Memory and RB- Memory, which were a mixture of CD27+/^lo^/-. RB+ 27+ Memory cells were further segregated in CD73+ and 73-subsets, just like naïve cells. A fifth memory population was uniquely defined by expression of CD95 (FasR), an activation molecule (Daniel and Krammer, 1994), and was primarily RB+ CD27+. The sixth memory subset was nonclassical, lacking uniform expression of both CD27 and CD45RB, but was CD19^hi^ CD11c+. Notably, this population also lacked expression of the chemokine receptors CD185 (CXCR5) and CD184 (CXCR4), suggesting these cells may not participate in germinal center responses (Cyster and Allen, 2019). There was some variation in subset size between donors, but all donors contained cells from all ten subsets across datasets assessed in this report (n=12, Figure 4C).

To assess single cell expression profiles, each subset was equally subsampled from donor-pooled cell observations, plotted by t-SNE (Van Der Maaten and Hinton, 2008), and colored by either subset or by molecule expression (Figure 4D). t-SNE was used in place of UMAP as UMAP resulted in excessive distances between populations, reducing resolution of local differences (Figure S2C). Subsets tended to form unique islands on the plot, providing an orthogonal validation of our classification methodology. Some cells seem to be mis-classified based on their t-SNE coordinates, such as several RB- Memory cells in the CD73+ Naïve island and CD95+ Memory cells in the Plasma island. However, the RB- Memory cells in question are IgG+ while naïve cells are, by definition, non-switched, and the CD95+ Memory cells are CD20+ and CD27-, while Plasma cells are CD20-CD27^hi^, demonstrating the accuracy of our classification approach (Maecker, 2012).

Transitional/naïve cells localized together into a single island, suggesting that these populations represent stages in a continuum from Transitional to CD73- Naïve to CD73+ Naïve cells. This continuum also follows a gradient of diminishing IgM and IgD, (*i.e.* IgM+ IgD+ Transitional, IgM^lo^ IgD+ CD73- Naïve, and IgM- IgD^lo^ CD73+ Naïve), despite the t-SNE plot being generated using only phenotypic, not isotypic molecules (Figure 4D). Notably, while there was a clear separation between transitional/naïve and memory subsets, no single molecule was sufficient to discriminate these cell types. Due to the uniqueness of their expression profile, plasma cells occupied a distinct island on the plot. While not completely homogenous, plasma cells were largely RB+, CD27^hi^, CD20-, CD38^hi^. As expected, these cells largely lacked surface Ig as assessed by surface light chain expression (see *Methods*, Figure S2D). IgA+ plasma cells and some IgM+ plasma cells were exceptions as they retained surface expression of Ig as previously reported (Pinto *et al*., 2013). We did not observe expression of surface IgG or IgD on plasma cells.

Three of the six memory populations organized into a distinct island suggesting these cells may also exist in a continuum: RB+ CD27- (CD73-) Memory to RB+ CD27+ CD73- Memory to RB+ CD27+ CD73+ Memory. Much like in the transitional/naïve continuum, differential isotype usage can be seen in this memory continuum with RB+ 27- Memory favoring IgD, RB+ 27+ 73- favoring IgM, and RB+ 27+ 73+ favoring IgG and IgA. RB- Memory was mostly class-switched, but had a more heterogenous profile, with varying levels of CD27, CD73, CD184, and CD183 (CXCR3), the chemokine receptor that facilitates homing to inflamed tissue (Kaminski *et al*., 2012). CD95+ cells also had heterogenous expression of several molecules, including CD11c and CD73, but tended to be class-switched, RB+, CD27+, and CD72-. CD19^hi^ CD11c+ cells formed a unique island characterized as CD20^hi^, CD38-, CD73-, CD40^lo^, and RB-. This subset had unique features based on isotype usage, with IgMD+ cells being CD27+/^lo^, CD185^lo^, CD183+ and CD95+/-, while class-switched cells were CD27-, CD185-, CD183-, and CD95-. We observed that several memory populations could also be further subdivided but were not further analyzed in this report: CD5+ IgMD+ RB+ CD27- CD73- CD38+ cells and CD9+ IgA+ RB+ CD27+ CD73- cells.

While IgH isotype was not used to meta-cluster cells, organizing B cells by phenotype resulted in an organization by isotype as well, in both pooled data (Figure 4E) and across individual donors (Figure S2E). Importantly, less than 0.26% of any transitional/naïve subset expressed a mature isotype. As class-switch recombination occurs only after activation, naïve cells are, by definition, not class-switched (Cyster and Allen, 2019). By canonical gating, 30% of IgG+ cells and 20% of IgA+ cells were left ungated due to an absence of CD27, demonstrating the insufficiency of CD27 alone as a memory molecule (Figure 4F, Figure S2F). In contrast, our meta-clustering approach correctly classified more than 99% of IgG+ and 98% of IgA+ cells as memory cells. Furthermore, by canonical gating, 45% of cells with an indeterminate isotype were left unannotated, as canonical gating of transitional and naïve cells relies on gating IgM+ and/or IgD+ cells that can often be absent or lowly expressed. No cells of indeterminate isotype were left unclassified by our meta-clustering approach, as it did not rely on binary definitions of molecule expression. An average of 10% of B cells across donors were of indeterminate isotype because of low/absent expression of surface Ig (Figure S2G) These cells are not mislabeled IgE+ cells as IgE+ B cells are exceedingly rare in healthy blood (Croote *et al*., 2018, Figure S2H). This is an important consideration in cytometry panel design as IgMD- is often used as a proxy for IgG+ or IgA+. Ig^lo^/- were found in all phenotypes and likely encompass a mixture of all isotypes. It is therefore essential to include probes against all four major isotypes if comparisons are to be made.

Beyond the associations established with our more comprehensively defined phenotypic subpopulations, Ig isotype usage by B cells is also known to affect downstream effector function and differentiation patterning (Dogan *et al*., 2009). As such, we organized B cells based on isotype alone and observed distinct patterns of expression of two key components of the B cell receptor (BCR) complex: surface Ig and CD79b (Figure 4G, Figure S2H). Surprisingly, the mature isotype, IgA, which is found exclusively on memory cells, had the lowest levels of expression of both molecules. Memory cells have a lower threshold for BCR-specific activation than naïve cells (Kurosaki, Kometani and Ise, 2015), despite IgA+ memory cells having much lower levels of receptor on their cell surface compared to IgM+ or IgD+ naïve cells. We also found that while CD79b and surface Ig expression were correlated in each isotype, IgG+ B cells had the highest correlation between surface Ig and CD79b expression (r=0.63, Pearson method), while IgA+ B cells had the lowest (r=0.32), suggesting IgA+ cells may not solely rely on CD79 for signaling, instead likely possessing a differential regulatory framework downstream of the BCR.

Given deep phenotypic heterogeneity within B cells in general, we asked whether phenotype or isotype contributed more to predicting expression levels of components of the BCR complex, surface Ig and CD79b. We created single-cell multiple linear regression models in which a cell’s phenotypic label (*e.g.* RB+ CD27- Memory) and isotypic label (*e.g.* IgA+) were used to predict the expression of either CD79b or surface Ig (see *Methods*, Figure 4H). While both were informative, a cell’s isotype contributed more than a cell’s phenotype for predicting the overall expression of the two molecules. Altogether these findings demonstrate that our high-dimensional classification organizes B cells into ten phenotypically distinct subsets with a more accurate partitioning of cells compared to canonical gating strategies. Additionally, these new phenotypic partitions displayed isotypic restriction that further contributed to a B cell’s identity, and predominantly dictated the expression of the BCR complex.

### Interrogation of B cell subset function reveals differential metabolic, biosynthesis, and immune signaling activity

To further investigate the functional relevance of our refined B cell classification scheme, we asked whether surface phenotypic diversity denoted differences in other underlying functional cell processes (Figure 5A). As metabolism has been shown to differ across T cell subsets (Buck, O’Sullivan and Pearce, 2015), as well as play a role in B cell activation (Boothby and Rickert, 2017), we interrogated single cell expression of key metabolic protein regulators across subsets. We stained healthy, human B cells from additional donors (n=3) with a mass cytometry panel targeting identified B cell surface molecules and metabolic enzymes (Figure 5A, Table S1). Cells were pooled from donors, clustered using FlowSOM, and assigned into the aforementioned B cell meta-clusters (see *Methods*, Figure 4).

**Figure 5:**
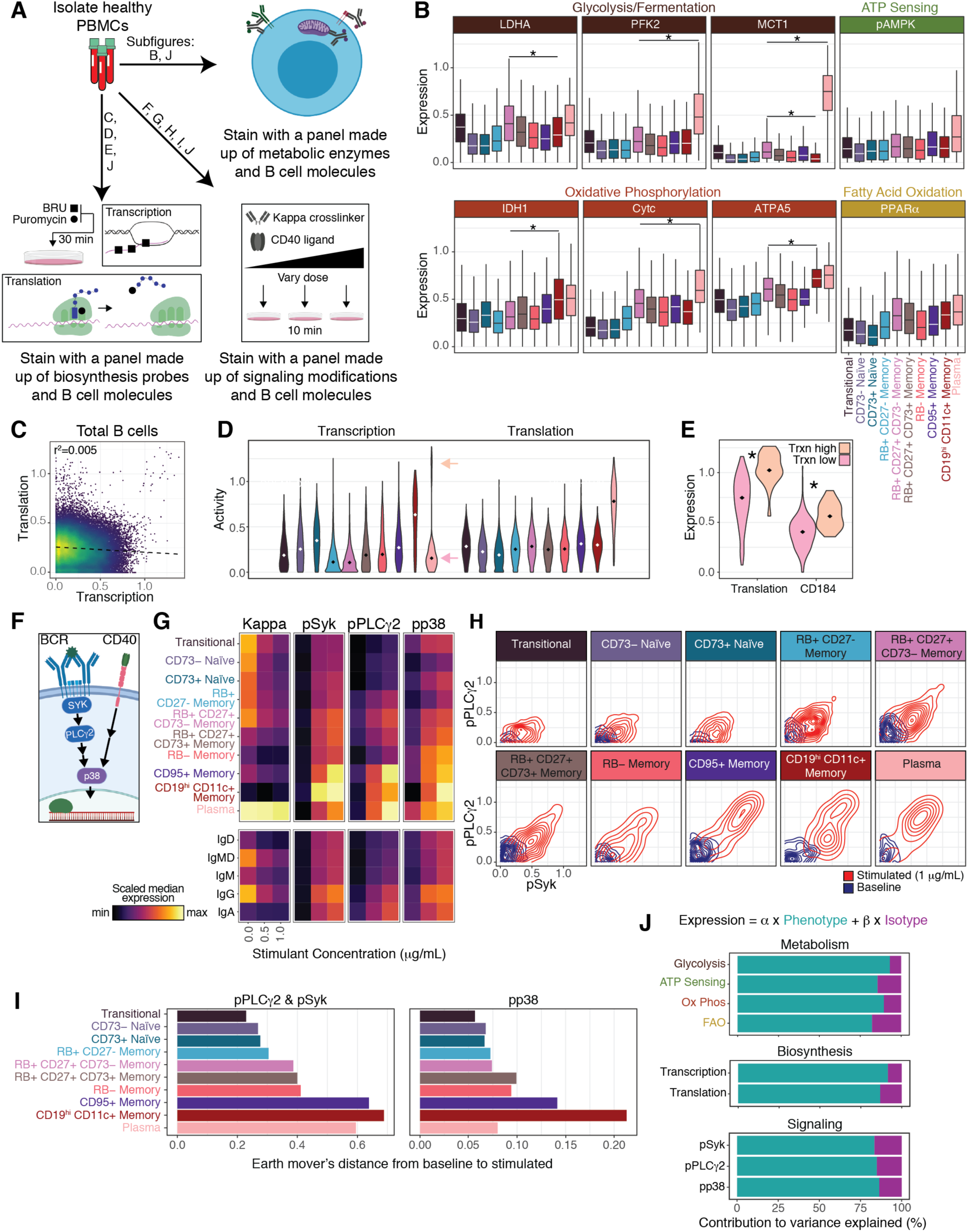
Interrogation of B cell subset function reveals differential metabolic, biosynthesis, and immune signaling activity. **A)** Experimental workflow for functional profiling. Peripheral blood was drawn from healthy, human donors (n=9), and density gradient centrifugation was performed. For metabolic profiling (n=3), cells were rested for 1 hour, fixed, and stained with a mass cytometry panel consisting of metabolic enzymes and B cell molecules. For profiling of RNA and protein biosynthesis activity (n=3), cells were rested for 30 minutes, then incubated with BRU and puromycin for an additional 30 minutes, fixed, and stained with a mass cytometry panel consisting of B cell molecules. For signaling profiling (n=3), cells were rested for 2 hours, stimulated with specified doses of anti-kappa and CD40L for ten minutes, fixed, and stained with a mass cytometry panel consisting of phosphorylated signaling molecules and B cell molecules. **B)** Boxplots of scaled expression of metabolic enzymes, segregated by donor-pooled B cell subset. Lines indicate median value, boxes indicate IQR, whiskers indicate first/third quartile +/- 1.5 IQR. Enzymes are colored by their associated metabolic pathway. Stars indicate significant differences in distribution between - subsets, p<0.005 by KS test with Bonferroni correction. **C)** Biaxial plot of donor-pooled total B cells, colored by density. Statistics and fitting line were calculated from a simple linear regression model. **D)** Violin plots of transcription and translation levels, segregated by B cell subset. Dots indicate median. Bimodal transcriptional activity in plasma cells is indicated by arrows. **E)** Violin plots of significantly differentially-expressed molecules between transcription^hi^ and transcription^lo^ plasma cells (p < 0.005, KS test with Bonferroni correction). Dots indicate median. Stars indicate significance. **F)** Cartoon depicting signaling pathways. After BCR crosslinking, SYK is phosphorylated, which induces phosphorylation of PLCγ2. Downstream of CD40-CD40L binding, p38 is phosphorylated and enters the nucleus to induce transcription. p38 is also induced by BCR crosslinking, to a lesser extent. **G)** Median expression of donor-pooled Igλ- B cells, grouped by phenotype (top boxes) or isotype (bottom boxes), separated by dose of stimulant (columns), and individually scaled by molecule. **H)** Biaxial density plots of signaling molecules separated by phenotype (boxes) of baseline (blue) and stimulated cells (1 μg/mL, red). **I)** Quantification of earth mover’s distance from baseline samples to stimulated samples (1 μg/mL) for pPLCγ2 and pSyk (left) and pp38 (right) **J)** Linear models were created to predict single-cell expression of metabolic pathways, biosynthesis activity, and cell signaling using IgH isotype class labels and meta-clustering population labels as predictors. The relative contribution of phenotype (cyan) and isotype (magenta) to the variance explained by each model was quantified. Glycolysis and oxidative phosphorylation contributions are reported as the mean contributions of their three associated enzymes.

To assess cellular metabolic predisposition, we quantified the expression of eight enzymes, associated with four metabolic pathways: glycolysis/fermentation, ATP sensing, oxidative phosphorylation (ox-phos), and fatty acid oxidation (Figure 5B). All subsets expressed all enzymes (e.g. transitional cells did not exclusively rely on fatty acid oxidation), but the levels of expression differed greatly by phenotype. Naïve cells had the lowest expression, which synergizes with the anergy we observed in their surface proteomes (Figure 2B-D), while RB+ CD27- Memory cells had an intermediate metabolic profile between naïve and memory subsets, much like their immune repertoire profiles (Figure 3G-H). Plasma cells had the highest median expression for all enzymes, and they were significantly greater than the next highest subset for phosphofructokinase 2 (PFK2) and monocarboxylate transporter 1 (MCT1), two glycolytic enzymes, and for cytochrome complex (CytC), part of the ox-phos pathway (p<0.005, KS test with Bonferroni correction). Within the memory compartment, RB+ CD27+ CD73- Memory and CD19^hi^ CD11c+ Memory cells had the highest median expression for all enzymes. Interestingly, CD19^hi^ CD11c+ Memory was significantly higher than RB+ 27+ 73- Memory for two ox-phos enzymes, isocitrate dehydrogenase 1 (IDH1) and ATP Synthase alpha subunit (ATP5A), while significantly lower for the glycolytic enzyme, MCT1 (p<0.005). These differences in pathway usage may be due to different functional roles, and therefore different metabolic needs.

In addition to single cell metabolic state, we recently developed an assay to quantify *de novo* RNA and protein synthesis in parallel with functional and phenotypic characteristics, by combining 5-Bromouridine (BRU) and puromycin labeling with mass cytometry (Kimmey *et al*., 2019). Interestingly, this study highlighted that B cells had the most variable and highest median transcriptional activity of all peripheral blood mononuclear cell types (PBMCs) with plasma cells being the most translationally active. We therefore asked if the B cell subsets identified here could reconcile the high variability we previously reported. To that end, we pulsed PBMCs from new donors (n=3) with BRU and puromycin, and stained cells with a B cell-centric mass cytometry panel to capture phenotype and functional reporters (Figure 5A, Table S1). Interestingly, across all B cells as a whole, *de novo* transcriptional activity explained very little of the variance observed in *de novo* translational activity (r^2^=0.005, Figure 5C), highlighting the differential regulation of these two processes.

To interrogate phenotypic associations, we broke donor-pooled B cells down into our identified subsets and plotted their synthesis activity (see *Methods,* Figure 5D). CD19^hi^ CD11c+ Memory cells had the highest median transcriptional activity, followed by CD73+ Naïve cells, which interestingly, had the lowest median translational activity. Given the anergy observed in the naïve department, it was surprising to see such a high level of transcriptional activity in these cells, and it is unclear what transcripts they are synthesizing, especially considering the low translational activity. As we previously reported, plasma cells had the highest median translational activity, but surprisingly, displayed bimodal transcriptional activity. We asked if any other molecules were differentially expressed between transcription^hi^ and transcription^lo^ plasma cells and found that translational activity and CD184 (CXCR4) expression were significantly higher in transcriptionally active plasma cells (p<0.005, Figure 5E). This transcriptionally active population may be long-lived plasma cells while the transcriptionally inactive population may be short-lived plasma cells. Long-lived plasma cells have been observed to upregulate CD184 to facilitate bone marrow homing and would require continuous transcriptional activity to facilitate constitutive Ig production and secretion (Nutt *et al*., 2015).

Given that transcriptional activity and translational activity were largely uncorrelated in total B cells, but transcription^hi^ plasma cells were more translationally active, we asked if the relationship between transcriptional activity and translational activity varied based on cell phenotype. We fit simple linear models to interrogate the relationship between transcriptional activity and translational activity in transitional/naïve clusters and separately in memory clusters (Figure S3A). Transcriptional activity and translational activity in transitional/naïve clusters had a strongly negative relationship (r^2^=0.61, p<1^−10^), but they had no relationship in memory cells (r^2^=0.02, p=1.32). Interestingly, transitional/naïve cells were generally more transcriptionally active, while memory cells were more translationally active. Total Ig levels (measured by intracellular staining) were correlated with translational activity in both transitional/naïve (r^2^=0.66, p<1^−12^) and memory (r^2^=0.28, p<1^−3^) clusters, but each regression had different coefficients and intercepts, so total Ig was only predictive of translational activity if the phenotypic subset was considered, highlighting the importance of proper subsetting in discovery and interpretation of biological findings and reinforcing the functionally-distinct nature of the subsets described here.

To assess if there were differences in immune activation sensitivity between B cell subsets, we interrogated cell signaling profiles in response to stimulation. PBMCs from new healthy donors (n=3) were incubated with varying doses of BCR crosslinker (anti-kappa light chain) and CD40 ligand (CD40L) for ten minutes, fixed, and stained with a B cell-centric mass cytometry panel that included antibodies against phosphorylated targets intrinsic to B cell signaling (Figure 5A, Table S1). We measured phosphorylation of spleen tyrosine kinase (pSYK) and the downstream phospholipase Cγ2 (pPLCγ2), two molecules involved in the signaling cascade caused by antigen recognition mediated by the BCR complex (Figure 5F, Kurosaki, Shinohara and Baba, 2010). We also measured phosphorylation of the stress-activated protein kinase p38 (pp38), which is strongly induced by CD40 stimulation, a molecule activated during antigen presentation to T cells, and weakly induced by BCR stimulation (Sutherland *et al*., 1996).

We segregated donor-pooled Igλ- B cells by our identified phenotypes, as well as by isotype, and assessed the median levels for each regulatory phosphorylation as a function of stimulant dose (Figure 5G). As expected, total kappa light chain diminished after stimulation as surface Ig was crosslinked, internalized, and degraded. Unsurprisingly, IgM+ and IgD+ cells had lower levels of signaling in response to stimulation, while cells with the mature isotypes, IgG and IgA, were the most potent responders. As we had seen in our previous datasets, IgA+ cells had the smallest quantity of Ig at baseline, yet responded with comparable potency to IgG+ cells, which had the highest quantity of Ig at baseline. This is particularly surprising as we had previously observed that IgA+ cells had the lowest expression of the BCR signaling molecule, CD79b. Segregating class-switched cells by phenotype, we found very little difference in signaling response between IgG+ and IgA+ cells of the same phenotype, despite their differences in BCR expression (Figure S3B). The only exception was plasma cells, in which IgG+ cells had weak responses compared to IgA+, probably due to a lack of surface Ig on IgG+ cells.

We then visualized the changes in phosphorylation state of the two molecules in the BCR complex signaling cascade, pSYK and pPLCγ2, on biaxial contour plots and found stark contrasts in distribution shifts between subsets (Figure 5H). While all populations responded to stimulation, only the plasma and memory subsets (particularly CD95+ Memory and CD19^hi^ CD11c+ Memory) contained a highly-responsive, double-positive population. To quantify the signaling response we calculated the earth mover’s distance between baseline and stimulated cells with earth mover’s distance and found that these two memory populations, along with plasma cells, were notably more responsive than all other subsets (Figure 5I, left, Figure S3C). Interestingly, the other seven populations self-organized from least to most responsive when we arranged subsets in a putative ordering of B cell maturation (Figure 4), suggesting that sensitivity to BCR-specific activation increases with this process.

Finally, we quantified the earth mover’s distance between baseline and stimulated cells for pp38, to assess CD40 signaling (Figure 5I, right, Figure S3D). While some of the trends were similar to the BCR complex signaling molecules (*e.g.* transitional/naïve cells were less responsive than memory cells), some differences emerged. Plasma cells had much lower signaling than in the BCR complex pathway, which is unsurprising as their primary function is antibody production, not T cell stimulation (Cyster and Allen, 2019). CD19^hi^ CD11c+ Memory signaling was uniquely high, despite having the lowest baseline expression of CD40, other than plasma cells (Figure 4B).

Given the heterogeneity of functional activity we observed across B cells, we asked if this variance was best explained by our phenotypic labels or by isotype. Using the same multiple linear regression approach used to quantify contributions to surface Ig and CD79b (Figure 4H), we quantified the relative contribution of phenotype and isotype usage to predict expression of metabolic pathway expression, biosynthesis activity, and signaling response (Figure 5J). In contrast to surface Ig and CD79b expression, a cell’s phenotype was much more informative than a cell’s isotype for predicting its metabolic, biosynthesis, and signaling profiles. Collectively, these findings demonstrate that unlike isotype alone, our new multiplexed phenotypic classification scheme for B cells captures functional distinctions in metabolic pathway usage, biosynthetic activity, and signal response to immune activation. Thus, these subset definitions represent a new framework to understand and assess the functional contributions of B cells to human immunity.

## Discussion

To interrogate deep phenotypic diversity in primary cells, we developed a highly-multiplexed single cell surface screening approach using mass cytometry and applied it to identify molecules that could segregate functional subsets of human B cells (Figure 1A). These methods are extensible to any cell population in a complex mixture and are particularly well-suited to rare cell types in limited specimens (*e.g.* clinical investigations) due to relatively low sample requirements. Furthermore, the approach is not confined to screening surface molecules and could easily be adapted and expanded to include intracellular targets as well. Our strict thresholding for considering a molecule positive greatly reduced the probability of false positives in our results (Figure 1C), but enriched for false negatives, so molecules found to be negative in the screen should not necessarily be seen as proof of their lack of expression in peripheral B cells. In fact, this dataset could be further mined for rarely and lowly expressed molecules.

Of 351 molecules screened, we identified 98 that were robustly expressed by peripheral blood B cells. Comparing molecules within canonical B cell populations, we identified 61 molecules that were differentially expressed between at least two populations (Figure 2A). We found a global downregulation of surface molecules in naïve cells coordinated with a specific upregulation of immunosuppressive molecules, suggesting these cells exist in an anergic state (Figure 2, 3B). Using an unsupervised approach, we then identified CD45RB as a new early classifier for memory cells and confirmed this proteomic finding with an orthogonal genomic analysis of IgH sequences within prospectively isolated B cells differentially baring CD45RB and the conventional memory marker CD27 (Figure 3). Koethe and colleagues previously demonstrated that the utility of the CD45RB antibody clone MEM55 (used in this study) to distinguish B cell subsets is based on differential glycosylation of CD45RB, rather than differential usage of the CD45RB isoform (Koethe *et al*., 2011). Unlike CD45RA and CD45RO splice isoforms in the switch of naïve to memory T cells, the switch of naïve to memory B cells identified here is accompanied by the post-translational modification of CD45RB, and therefore impossible to detect with mRNA sequencing, instead requiring a proteomic readout.

We found that RB+ CD27- B cells had IgH mutation rates greater than RB- CD27- naïve cells and lesser than RB+/- CD27+ memory populations, suggesting they form an early memory population. This complements a finding reported in human tonsil by Jackson and colleagues in which RB+ naïve B cells (defined as IgD+ CD27- CD38-) had a significantly higher mutational burden in IgV_H_4 IgM transcripts than RB- naïve B cells (Jackson *et al*., 2009). Interestingly, in that report, within germinal center B cells (defined as IgD- CD38+), RB+ cells had lower mutational burdens than RB- cells. In our own dataset, within canonical memory populations (CD27+), RB+ cells had no significant difference in mutational burden than RB- cells, though the RB- population had greater sequence diversity. This greater diversity may be indicative of a longer-lived memory population which provides protection against a broad range of infectious agents, experienced over a longer span of time. Within clonal lineages, there was significant coordination of expression of CD45RB and CD27, mirroring the coordination of isotype usage seen within clonal lineages by Horns and colleagues (Horns *et al*., 2016).

Based on our quantification of B cell phenotype, isotype usage, metabolism, biosynthesis activity, and signaling, we proposed a new classification scheme for peripheral blood B cells (Figure 6A). The scheme only requires seven molecules (including CD19) to differentiate all ten B cell subsets, as opposed to the canonical gating approach which required six molecules to differentiate five populations (Figure 1A-B). Furthermore, using the canonical approach, several functionally-distinct subsets are either A) not segregated from other populations (*e.g.* hyper-responsive CD95+ Memory classified with all other switched memory cells), B) entirely misclassified (*e.g.* RB+ 27- Memory classified as naïve cells), or C) spread amongst several subsets (*e.g.* CD19^hi^ CD11c+ Memory cells spread across ungated, naïve, non-switched memory, and switched memory gates in the conventional scheme).

**Figure 6:**
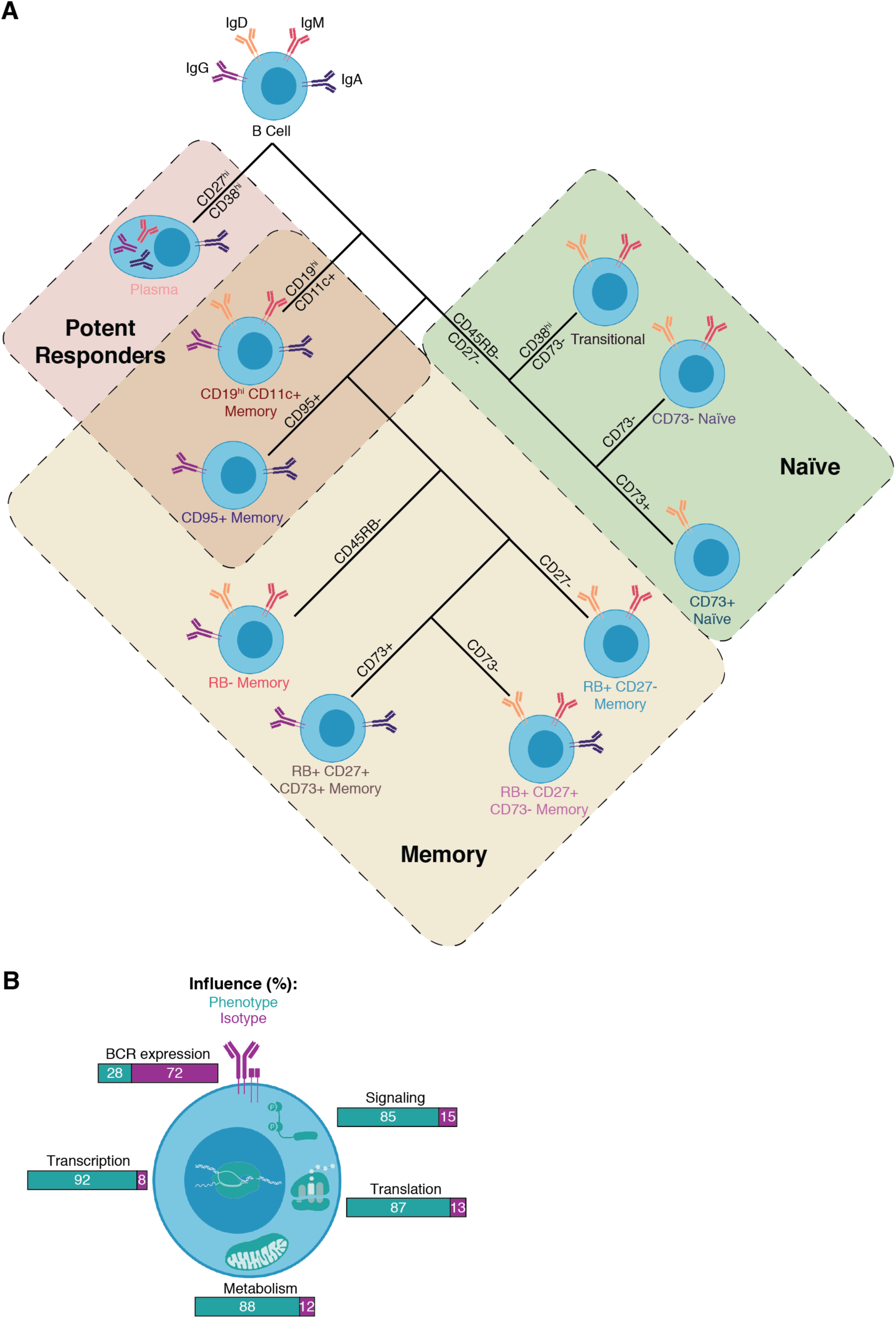
A new classification scheme for peripheral B cells and a summary of the contributions of phenotype and isotype to cell activity. **A)** Schematic overview of proposed classification scheme. Each subset is labeled with isotypes present in at least 10% of cells in that subset (colored Ig). Subsets are putatively ordered from least mature to most mature beginning with Transitional cells and moving clockwise to RB- Memory. **B)** Summary of relative contributions of isotype and phenotype to various cell activities. Quantification of metabolism and signaling are reported as means of the contributions of all pathways (metabolism) or all phospho-molecules (signaling).

While the initial discovery of our ten populations relied on hierarchical meta-clustering of over-clustered data (Figure 4), subsequent identification of these populations relied on a simple gating scheme of over-clustered data (Figure 5). The initial over-clustering of high-dimensional data takes advantage of all measured parameters (quantification of 40+ molecules) to facilitate easy grouping of phenotypically-similar cells, while the manual gating of clusters ensures consistent classification from dataset to dataset. Importantly, these populations can also be manually gated without clustering (*e.g.* for prospective isolation by FACS), though the purity and yield would suffer using binary gating strategies rather than high-dimensional cell properties (Figure S2B).

We subdivided antigen-inexperienced cells (RB- CD27-) into three populations which seem to form a continuum of maturation from Transitional to CD73- Naïve to CD73+ Naïve. We found there was a reduction in translational activity and increase in transcriptional activity through this maturation process – future studies could address which transcripts are being generated by CD73+ Naïve cells. While CD73+ Naïve cells had seemingly more anergic profiles (IgD^lo^ IgM- CD79b^lo^/-) than CD73- Naïve cells (IgD+ IgM^lo^ CD79b+/^lo^), both had similar signaling responses to stimulation (Figure 4B, 5G-I). Similarly, transitional cells had the highest median quantities of IgM, IgD, and CD79b for any B cell subset (Figure 4B) but had the weakest response to stimulation (Figure 5G-I), demonstrating that BCR quantity is a poor predictor of activation potency, even in cells of the same isotype.

While plasma cells formed a single subset that largely overlapped with canonical definitions (CD27^hi^ CD38^hi^ CD20-), we found bimodal transcriptional activity, where the transcription^hi^ population had significantly higher expression of the chemokine receptor, CD184 and higher translational activity (Figure 5D-E). While these two populations are largely similar in phenotype, these differences may distinguish long-lived (CD184^hi^ translation+++ transcription^hi^) from short-lived (CD184+ translation++ transcription^lo^) plasma cells. This highlights the utility of quantifying intrinsic cellular biochemical activity, such as molecule biosynthesis, as a readout for functionally distinct cell subsets and advocates for further studies like this on plasma cells in the context of immune challenge.

We segregated memory cells into six distinct populations. RB+ CD27- Memory cells formed an early memory population that almost exclusively displayed the immature isotypes, IgM and IgD (Figure 4E). RB+ CD27+ CD73- Memory cells also tended to use the immature isotypes, but demonstrated a greater bias towards IgM over IgD, while RB+ CD27+ CD73+ were typically class-switched. These three populations seemed to form a continuum of maturation (RB+ CD27- Memory to RB+ CD27+ CD73- Memory to RB+ CD27+ CD73+ Memory) not only based on isotype usage, but also due to increasing levels of CD82, CD183, CD29, CD40, and HLA-ABC (*i.e.* MHC class I), and decreasing levels of CD1c, CD305, and CD72 (Figure 4B, D). RB- Memory was also predominantly class-switched, but had a more heterogenous phenotypic profile, with varying levels of CD73, CD183, and CD184 (Figure 4B-E). This phenotypic heterogeneity mirrored these cells’ IgH sequence diversity (Figure 3H), and along with the high mutation rates (Figure 3G) and mature isotype usage, may represent a long-lived memory population that mediates protection against a lifetime of pathogens. This population may therefore harbor unique, protective monoclonal antibodies that could have relevance in vaccine design and as therapeutics.

CD95+ Memory was highly responsive to stimulation, particularly in the BCR signaling pathway (Figure 5G-I). These cells were mostly RB+, 27+, CD184-, and class-switched, but had mixed expression of the inflammatory chemokine receptor, CD183, the integrin, CD11c, and the immunoregulatory ecto-nucleotidase, CD73 (Figure 4B, D). Along with plasma cells, CD95+ Memory are the only B cells in our data that lacked expression of CD72, a negative regulator of B cell activation, which may in part explain their high sensitivity to BCR stimulation. CD95 can be expressed on activated lymphocytes as a mechanism for limiting inflammation, as ligation of CD95 can lead to apoptosis (Daniel and Krammer, 1994), so CD95+ Memory cells may simply be short-lived effector cells. The ligation of CD95, however, does not exclusively result in apoptosis and it may be relevant in lymphocyte activation, survival, and proliferation (Peter *et al*., 2007), so rather than accelerating cell death, CD95 may be endowing these cells with increased potency and longevity. The presence of this cell type in all twelve healthy blood donors (Figure 4C) suggests that this is a more permanent population, though future studies could address this question.

We also identified a CD19^hi^ CD11c+ Memory population that shares some common features with several populations that have been described in the context of autoimmunity, infection, and aging (Karnell *et al*., 2017). Our population is CD20^hi^, RB-, CD21^lo^/-, CD73-, CD38-, CD22^hi^, CD24-, CD62L-, CD35-, CD268^lo^, CD44^lo^, HLA-ABC^hi^, and a mixture of CD27+ non-switched cells and CD27- switched cells (Figure 4B, D). The population had high expression of metabolic enzymes, biased towards aerobic oxidative phosphorylation over more anaerobic glycolysis (Figure 5B). CD19^hi^ CD11c+ Memory cells had the highest median transcriptional activity of all B cell subsets and median translational activity only lower than the other two potent responders to activation: CD95+ Memory and Plasma cells (Figure 5D). These cells lack CD184, have low expression of CD40, and mixed expression of CD185, suggesting they do not participate in germinal center reactions, and yet they had the most potent response to activation in both the BCR and CD40 signaling pathways (Figure 4B, 5I). This contrasts previous reports that have indicated cells with similar phenotypes are hyporesponsive to BCR and CD40 activation (Isnardi *et al*., 2010, Rakhmanov *et al*., 2009). This discrepancy may be explained by several factors. Our stimulation was only ten minutes long and quantified downstream signaling events, while previous studies used multi-day stimulations with different readouts (*e.g.* proliferation and upregulation of CD69). Our population was segregated by high-dimensional clustering, not binary gating, so the cells included and excluded by the classification is likely different (*e.g.* our population is CD27+/-, while cells in other reports are gated CD27-). We stimulated the BCR with anti-kappa light chain and then analyzed only Igλ- cells. In previous studies, BCR was stimulated with anti-IgM, but this approach may bias results as our population was <50% IgM+. CD11c expression can be induced in B cells after activation and therefore this population may simply be recently activated B cells (Postigo *et al*., 1991), but the presence of this cell type in all twelve healthy blood donors (Figure 4C) and its unique functional profile suggests an alternate role.

In this study, we also evaluated the relative contribution of phenotype and isotype to predicting various quantifications of cellular processes (Figure 4H, 5J, 6B). We found that isotype was more informative than phenotype for predicting the overall expression of the BCR complex. Given that intracellular signaling domains differ between the isotypes (Martin and Goodnow, 2002), this is not entirely unexpected, as more sensitive isotypes would require less surface Ig for activation. For all other processes measured (transcriptional activity, translational activity, metabolic profile, and cell signaling), phenotype was the more informative predictor. This is unsurprising as our phenotypic organization of the B cell compartment organized cell by isotype as well. Furthermore, antibody isotype describes the state of a single gene locus (IgH) while phenotype can refer to any number of proteins expressed in the B cell proteome. Even differences in signaling that are mediated by differential isotype usage would only result in differential functional outcomes when mediated by other molecules. While isotype usage captures important information about a cell and should not be disregarded, our dataset suggests that a combination of the correct phenotypic molecules better segregates B cells into functional groups, as it has been established for other immune effector cells as well.

Here, our deep phenotypic profiling with multi-omic integration of numerous single cell functional readouts on peripheral blood in healthy individuals reveals the identity of new, more granular populations comprehensively mapping B cell identity in the human. The quantitative assessment of the contribution of phenotype (*i.e.* cell identity) versus isotype usage across several cellular processes highlights the need for analysis beyond repertoire sequencing and isotype identity for understanding human B cell immune function. While we only interrogated peripheral blood, observation of these populations and molecules by high-dimensional imaging (Keren *et al*., 2018) in secondary lymphoid organs may provide new insights into germinal center processes. Our findings should serve as a resource for future studies investigating human B cell immunity in the context of disease, as several populations and/or molecules described here may be crucial to understanding the pathogenesis or protection it confers.

## Methods

### Cell processing

Deidentified human blood was obtained from healthy adult donors under informed consent (Stanford Blood Center). Use of these samples was approved by Stanford’s Institutional Review Board. Peripheral blood mononuclear cells (PBMCs) were isolated from Trima Accel leukocyte reduction system (LRS) chambers (Terumo BCT) or heparinized tubes using Ficoll-Paque Plus (GE Healthcare) density gradient centrifugation according to the manufacturer’s instructions. For long-term storage (surface screen only), PBMCs were resuspended in FBS with 10% DMSO and stored in liquid nitrogen at a density of 1-5 × 10^7^ cells/mL. Cryopreserved PBMCs were thawed into cell culture medium (CCM; RPMI 1640 containing 10% FBS, and 1× L-glutamine; Thermo Fisher Scientific) supplemented with 25 U/mL benzonase (Sigma-Aldrich). and pelleted for 5 min at 250 *g*. Where indicated, cells underwent magnetic lineage depletion according to the manufacturer’s instructions using BD Streptavidin Particles Plus and the BD IMag Cell Separation Magnet (BD Biosciences) with biotinylated anti-CD3 (surface screen samples) or a cocktail of biotinylated antibodies consisting of CD3, CD7, CD15, CD33, CD56, CD61, and CD235ab (other samples). The biotinylated antibody cocktail was detected by labeled anti-biotin (CyTOF) and streptavidin (FACS) and further depleted *in silico*.

### Metabolism, biosynthesis activity, and stimulation assays

PBMCs were rested in 37 °C 5% CO_2_ incubator for 30 min (biosynthesis activity assays), 1 hour (metabolism assays), or 2 hours (stimulation assays) at 5 x 10^6^ cells/mL in CCM. Metabolism samples were fixed in 1.6% PFA in PBS for 10 min and then barcoded as previously described (Zunder *et al*., 2015). After rest, biosynthesis activity assay samples were spiked with 2 mM BRU and 10 μg/mL puromycin and incubated for an additional 30 min (Kimmey *et al*. 2019) and then fixed and barcoded. Stimulation samples were resuspended in CCM with 3.3 mM H_2_O_2_ (Irish *et al*., 2010) and specified doses of anti-kappa F(ab’)_2_ (Southern Biotech) and CD40L (BioLegend) for 10 min and then fixed and barcoded.

### CyTOF antibody conjugation, staining, and data acquisition

Antibody conjugation, staining, and data acquisition were performed as previously described (Hartmann, Simonds and Bendall, 2018). Briefly, metal-isotope labeled antibodies used in this study were conjugated using the MaxPar X8 Antibody Labeling kit per manufacturer instruction (Fluidigm), or were purchased from Fluidigm pre-conjugated. Each conjugated antibody was quality checked and titrated to optimal staining concentration using a combination of primary human cells and/or cancer cell lines (Figure S1C). Cells were suspended in TruStain FC blocker for 10 minutes at RT and washed in cell staining media (CSM: PBS with 0.5% BSA and 0.02% sodium azide and benzonase 25×10^8^ U/mL (all Sigma)) prior to staining. All surface staining was performed in CSM for 30 min at RT. Cells were washed in CSM and resuspended in cisplatin for 5 minutes to label non-viable cells (Sigma, 0.5 μM final concentration in PBS). Cells were washed in CSM and fixed with 1.6% PFA in PBS for 10 min at RT and (if intracellularly stained) washed in CSM and permeabilized with MeOH for 10 min on ice. Intracellular and anti-biotin staining was performed in CSM for one hour at RT. Before acquisition, samples were washed in CSM and resuspended in intercalation solution (1.6% PFA in PBS, 0.02% saponin (Sigma) and 0.5 μM iridium-intercalator Fluidigm)) for 1 h at RT or overnight at 4 °C. Before acquisition, samples were washed once in CSM and twice in ddH_2_O. All samples were filtered through a cell strainer (Falcon), resuspended at 1 × 10^6^ cells/mL in ddH_2_O supplemented with 1x EQ four element calibration beads (Fluidigm), and acquired on a CyTOF2 mass cytometer (Fluidigm). Barcoded samples were acquired using the Super Sampler injection system (Victorian Airship).

### FACS sorting, gDNA extraction, and immune repertoire sequencing

PBMCs were lineage depleted and Fc blocked as described above. Cells were surface stained in CSM in the dark for 30 minutes on ice and then washed in CSM. Prior to data acquisition, cell suspensions were spiked with 7-AAD to label non-viable cells. Non-transitional/non-plasma B cells were gated as singlet, viable, CD45+, lin-, CD19+, CD38lo/- and then sorted from the four quadrants of the CD27 x CD45RB biaxial plot using a BD FACS Aria II (BD Biosciences). Approximately 100,000 cells per subset per donor were sorted into tubes. Cells were lysed and gDNA was extracted using the QIAamp DNA Micro kit (Qiagen) according to the manufacturer’s instructions. DNA was frozen and shipped to Adaptive Biotechnologies for IgH library preparation and next-generation sequencing using the immunoSEQ Assay (Adaptive).

### CyTOF data pre-processing

Acquired samples were bead-normalized using MATLAB-based software as previously described (Finck *et al*., 2013). Where applicable, barcoded data was debarcoded using MATLAB-based software (Zunder *et al*., 2015). Normalized data was then uploaded onto the Cytobank analysis platform for gating (Figure 1B, S1A; Kotecha, Krutzik and Irish, 2010). Gated data was downloaded and further processed with the R programming language (http://www.r-project.org) and Bioconductor (http://www.bioconductor.org) software. Data was transformed with an inverse hyperbolic sine (asinh) transformation with a cofactor of 5. Molecules on the screen compromised by bleed from other channels or by any other technical considerations were removed. Conserved molecules from the surface screen were quantile normalized by donor to correct for technical variation between CyTOF runs. Peak normalization (alignment of mode of positive population) of each molecule was applied to normalize samples from different donors in Figure 4 only, as those samples were not barcoded and stained in the same tube (Figure S2A). For all experiments, each molecule was scaled to the 99.9^th^ percentile of expression of all cells in that experiment for comparability between parameters. Individual cell quantifications from all donors were pooled together for all analyses except where noted to utilize all observations acquired.

### Immune repertoire sequencing pre-processing

Templates per sequence (number of unique cells with identical IgH sequences) was determined by Adaptive Biotechnology based on number of sequencing reads normalized to spiked-in controls of “artificial” IgH sequences. Sequences with less than ten reads were eliminated from the analysis. The Immcantation pipeline (Vander Heiden *et al*., 2014; Gupta *et al*., 2015) was used for downstream processing and analysis. V and J gene usage was determined using IgBlast (Ye *et al*., 2013) on the IMGT database (Lefranc *et al*., 2015) and corrected by Bayesian inference of each donor’s genotype. Clonal lineages assignments of sequences were made with the following requirements: same donor, identical V and J gene usage, identical CDR3 length, and a hamming distance to another member of the lineage beneath the set threshold. This distance threshold was determined for each donor by fitting a generalized mixture model based on the density plot of the hamming distance of the CDR3 to its nearest neighbor for each sequence. This creates a bimodal distribution of sequences with a clonal relative (lower peak) and sequences without a clonal relative (higher peak). The intersection of the two fitted gaussians was set as the distance threshold for clonal membership. Germline sequences were inferred for each clonal lineage and silent and non-silent mutations outside of the CDR3 were quantified as deviations from the inferred germline.

## Data Analysis

### GO quantifications

“Biological Processes” gene ontology annotations for all molecules on the surface screen were compiled from UniProt (https://www.uniprot.org). This resulted in over 2,000 unique annotations, so GO terms were collapsed into 30 parent terms using the Generic GO Term Mapper (Figure S1B, https://go.princeton.edu/cgi-bin/GOTermMapper).

### Surface screen thresholding

To avoid subjectivity in determining whether each molecule was present or absent on B cells, we set a uniform threshold for positivity, mandating that the 99.9^th^ percentile of expression in B cells for each molecule was at least 40 raw counts (2.78 asinh-transformed value) to be considered positive. The cutoff was set at a high value to prevent inclusion of false positives with high background staining, at the expense of enriching for false negatives. This approach provided confidence that all molecules assessed as positive were truly present on B cells, while those assessed as negative were either absent on B cells or were present at low levels. The 99.9^th^ percentile was used instead of median values for thresholding, in order to capture molecules expressed by a small subset of B cells (e.g. IgA) and not just those uniformly expressed by all B cells (e.g. beta-2 microglobulin, Figure 1C). Furthermore, with a mean of 245 thousand B cells assessed with each CyTOF panel in the screen, a molecule was on average only considered positive if ∼245 cells express the target at a high level.

### Ig quantification

To quantify surface Ig (surface stain, Figure 4B, D, G) and total Ig (intracellular stain, Figure S3A) across IgH isotypes, Igκ and Igλ light chains were independently measured on two different CyTOF channels. The two distributions were peak normalized within each experiment (Figure S2D) and then each cell was given a new parameter, surface or total Ig, calculated as the pairwise maximum of the two peak normalized light chain channels. As B cells can only express a single light chain isotype, the pairwise maximum represents true signal, while the pairwise minimum represents noise and can therefore be discarded. This approach provides an independent quantification of Ig expression for single cells that is not biased by antibodies with differing affinities to the various IgH isotypes.

### Dimensionality Reduction

To visualize co-expression of molecules measured on different cells in the surface screen, cells stained with different CyTOF panels were plotted together on a single UMAP plot using the umap package in R (Figure 3A-B). 2,500 cells from each of the 12 CyTOF panels were randomly subsampled and used to generate a UMAP plot based on the expression of molecules positive on B cells that are conserved in all panels: CD45, CD19, CD24, CD38, CD27, IgM, and IgD. This ensured that cells of similar phenotype localized into similar coordinates, facilitating qualitative assessment of molecule co-expression between molecules that were not measured on the same cell. Color overlay of molecule expression for a given molecule used only cells from the screen on which that molecule was measured for the visualization. For visualization of surface molecule expression of our meta-clusters, 1,000 cells were randomly subsampled from each of the ten B cell subsets and used to generate a t-SNE plot with the Rtsne package in R (Figure 4D) or a UMAP plot with the umap package in R (Figure S2C). The plot was generated based on expression of CD32, HLA-ABC, CD185, CD5, CD72, CD45RB, CD183, CD81, CD21, CD29, CD79b, CD20, CD40, CD268, CD11c, CD24, CD95, CD19, CD27, CD73, CD62L, CD99, CD44, CD38, CD184, CD82, CD305, CD22, and surface Ig. Subsampling by B cell subset facilitated visualization of heterogeneity within and between populations without the map being dominated by the most abundant populations. Isotype was not used to generate the map to prevent artificial separation of phenotypically similar cells.

### Immune repertoire analyses

Mutation frequency was calculated as the frequency of mutations from the reconstructed germline sequence to input sequence (Figure 3G). The D region and N/P nucleotides were excluded from the calculation as they are difficult to accurately call and reconstruct. Mutations were binned as either silent (no amino acid change) or replacement (amino acid change). Sequence diversity was calculated using the general form of the diversity index (Hill, 1973) over a range of orders (Figure 3H). 95% confidence intervals were generated by 200 bootstrap resampling calculations, as previously described (Gupta *et al*., 2015). For clonal analysis, all sequences were labeled by their population of origin and then these labels were then randomly permuted (Figure 3I). For each population label, the frequency in which a sequence from population X shared a clonal lineage with a sequence from population Y was quantified and then repeated for all combinations of the four populations. This calculation is asymmetric as the frequency in which a sequence from population X shares a lineage with a sequence from population Y is not the same as the frequency in which a sequence from population Y shares a lineage with a sequence from population X. This process was repeated 200 times to create a null distribution and then z-scores of the frequencies were derived for the observed data using the original population labels (Figure 3J).

### Clustering

For initial subset discovery (Figure 4) cells were over-clustered into 169 clusters using FlowSOM with all molecules as input. Clusters were then hierarchically clustered as either “antigen-inexperienced” or “antigen-experienced” based on median expression of CD45RB, CD27, CD305, CD44, and CD11c. Antigen-inexperienced clusters were then hierarchically clustered into three subsets: (1) Transitional, (2) CD73- Naïve, and (3) CD73+ Naïve, based on expression of CD38, CD79b, and CD73. Antigen-experienced clusters were hierarchically clustered into four subsets: (1) Plasma, (2) CD95+ Memory, (3) CD19hi CD11c+ Memory, and (4) other memory based on expression of CD20, CD268, CD95, and CD11c. Other memory clusters were hierarchically clustered into two subsets: (1) RB- Memory and (2) RB+ Memory based on expression of CD45RB. RB+ Memory was then hierarchically clustered into three subsets: (1) CD27- Memory, (2) RB+ CD27+ CD73- Memory, and (3) RB+ CD27+ CD73+ Memory based on expression of CD27 and CD73. This entire procedure resulted in the identification of ten unique subsets. This approach was selected for several reasons: A) The initial over-clustering step groups phenotypically similar cells based on high-dimensional data and prevents arbitrarily drawing lines between overlapping populations based on a single molecule, as in canonical gating schemes. B) Hierarchical clustering allows segregation of cells into large groups (antigen-inexperienced vs antigen-experienced) before further subsetting. This mirrors canonical gating, where T cells are segregated from B cells before expression of CD4 or CD8 is considered. C) This approach allows us to use specific molecules to subset specific groups of cells. Once we have determined that expression of a parameter is uniform in a group (e.g. CD45RB in naïve cells), the use of that molecule as a clustering parameter will only add noise to the model. Instead, we selected molecules for each group where there was meaningful differential expression (e.g. CD73 in naïve cells).

For subsequent subsetting of other datasets (Figure 5 & 6), cells were over-clustered with surface and intracellular Ig molecules and then manually segregated into the ten subsets in a gating scheme similar to Figure S2B. This ensured consistency of classification between datasets. For signaling data, only unstimulated cells were clustered and classified so that Ig levels could be used in the initial clustering step. After crosslinking, Ig levels diminish, so clustering based on these molecules causes a segregation of cells based on stimulation dose. Stimulated cells were instead classified into subsets based on a KNN (k=3) classifier trained on unstimulated data, using only Euclidean distance of surface molecule expression, which is not altered by the short stimulation.

### Statistics

All statistical tests for differences in distribution of molecules between B cell subsets from mass cytometry was performed on equally subsampled populations using the KS test with Bonferroni correction, and only considered significant if p < 0.005. This test is sensitive to both changes in mean and shape of a distribution, and can therefore detect if even a fraction of a subset has a change in expression compared to the reference population. For comparisons of mutation frequencies between sorted B cell populations (Figure 3G), the Wilcoxon rank sum test was performed with Bonferroni correction, and only considered significant if p < 0.005. To determine the contribution of phenotype and isotype to various processes (Figure 4H, 5J), a multiple linear regression model was used for each response. Each observation (cell) was labeled with two discrete predictors: phenotype (B cell subset membership determined by clustering) and isotype (determined by manual gating). These two predictors were used to regress the continuous expression value of the desired response variable (e.g. CD79b expression). The relative contribution to variance explained by each predictor was calculated using the “lmg” metric of the relaimpo R package (Grömping, 2015).

### Figure generation

Figures were prepared using BioRender (http://www.biorender.com) and Illustrator (Adobe).

## Acknowledgements

Thanks to Leeat Keren and Erin McCaffrey for discussion and comments. Thanks to Jennifer Wang, Trevor Bruce, and the Stanford Blood Center Flow Cytometry Core for experimental assistance. Thanks to the Stanford Immunology Graduate Program leadership and administration for training and support. D.R.G. was supported by a Stanford Graduate Fellowship and a Bio-X Stanford Interdisciplinary Graduate Fellowship. A.G.T. was supported by a Damon Runyon Cancer Research Foundation – DRCRF (DRG-118-16) and Stanford Department of Pathology Seed Grant. J.P.O. was supported by a Canadian Institute of Health Research Postdoctoral Fellowship. F.J.H was supported by the EMBO organization (EMBO Long-Term Fellowship), the Novartis Foundation for medical-biological Research and the Swiss National Science Foundation (SNF Early Postdoc Mobility). S.C.K. was supported by the NIH/NIGMS Cell and Molecular Biology Training Grant (T32GM007276). A.A.C. was supported by the NIAID of the National Institutes of Health under award number 5T32AI007290-32 and the National Science Foundation Graduate Research Fellowship Program under Grant No. DGE – 1656518. S.C.B. was supported by the DRCRF Fellowship (DRG-2017-09), the NIH 1DP2OD022550-01, 1R01AG056287–01, 1R01AG057915-01, 1-R00-GM104148-01, 1U24CA224309-01, 5U19AI116484-02, U19 AI104209, The Bill and Melinda Gates Foundation, and a Translational Research Award from the Stanford Cancer Institute

## Author contributions

D.R.G, A.G.T., and S.C.B. conceived and designed the study. D.R.G., A.G.T., J.P.O., F.J.H., S.C.K., A.A.C., and L.B. performed experiments. D.R.G. analyzed data and wrote the manuscript. S.C.B. supervised and funded the study.

## Code and data availability

All code and data will be made publicly available at the time of publication.

---

**Figure S1:**
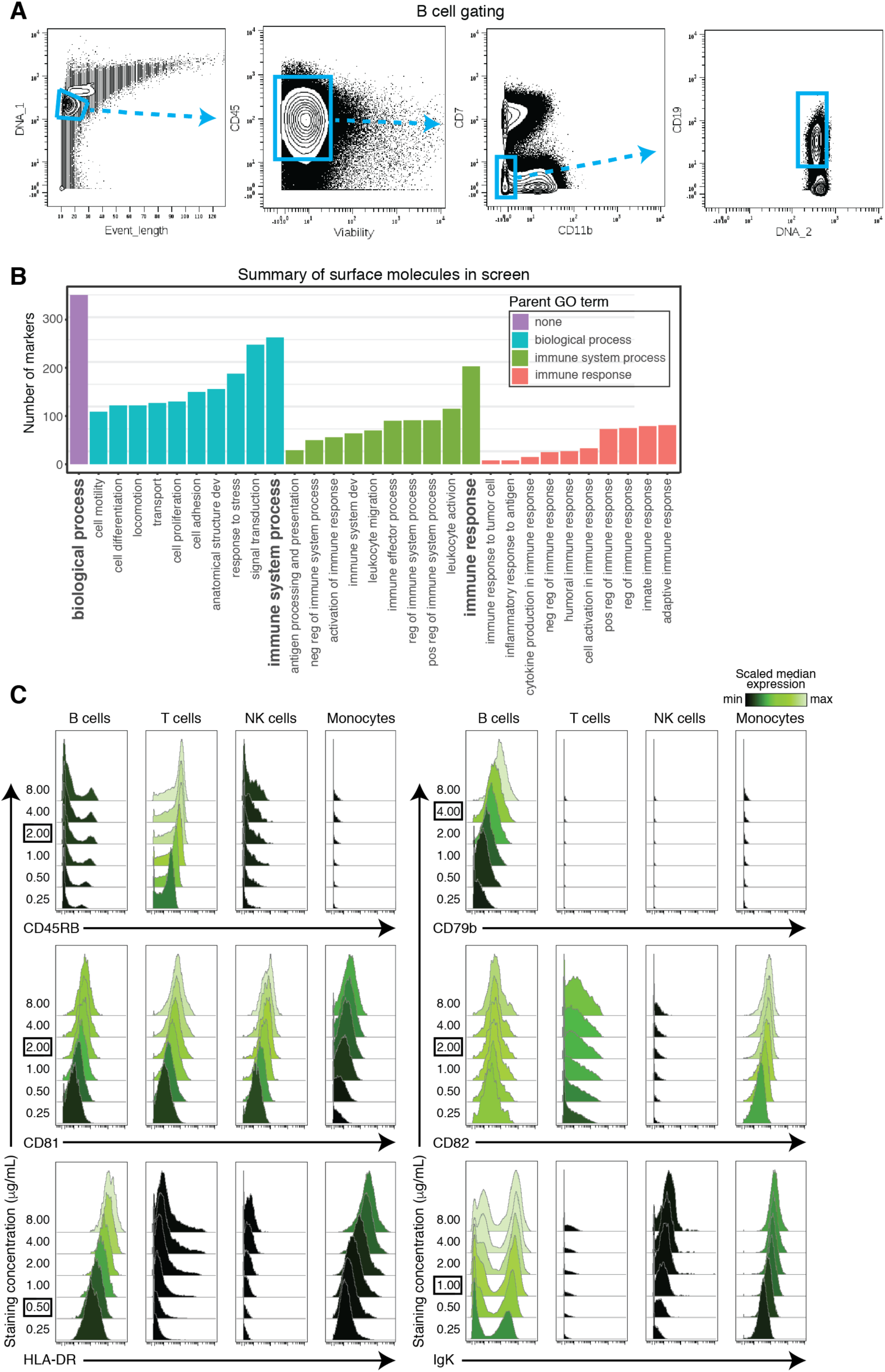
Surface screen gating, summary, and quality control - related to Figure 1. **A)** Representative plots from one donor of the gating strategy for total B cells in the surface screen. **B)** Quantification of the number of surface molecules in the surface screen that were associated with specific GO annotations. Parent terms are bolded. **C)** Representative plots of surface screen antibody staining concentration titrations of healthy PBMCs, arranged by immune population (columns) and staining concentration (rows, μg/mL). Staining concentration used in the screen is boxed. Color represents scaled median intensity of each molecule. All populations are defined as CD45+ lin-. Additionally, B cells are CD19+, T cells are CD3+, NK cells are CD56+, and monocytes are CD14+.

**Figure S2:**
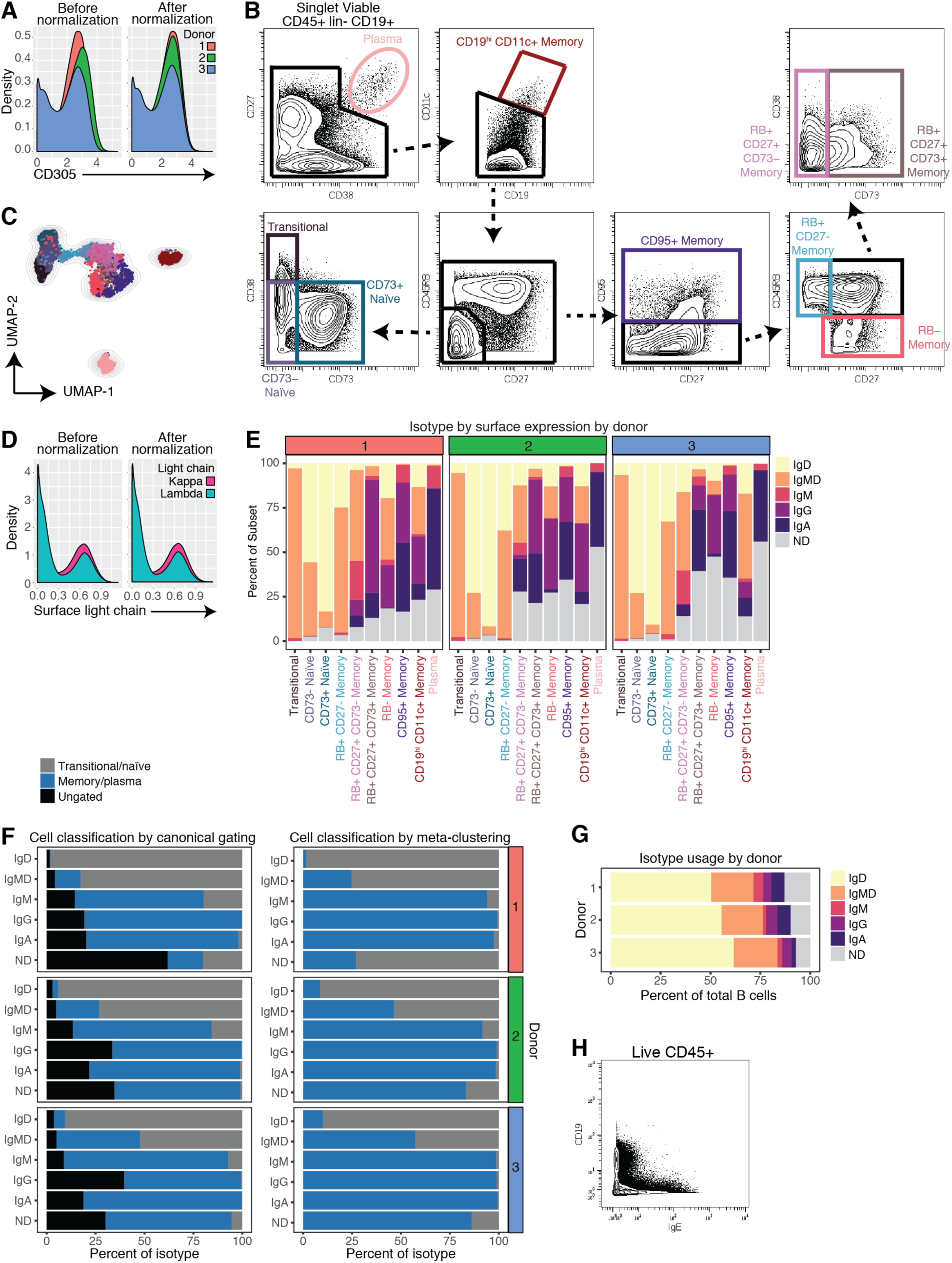
Quality control, data processing, and individual donor contributions to phenotypic characterization - related to Figure 4. **A)** Representative density plots of asinh-transformed expression value of CD305 for total B cells of three donors (colors) before and after peak normalization. **B)** Representative plots from a single donor of the gating strategy for B cell subsets. **C)** UMAP plot generated using the same input as the t-SNE plot in Figure 4D - an equal subsampling of 1000 cells from each B cell subset using only phenotypic molecules. Cells are colored by subset identity as in B). **D)** Representative density plots of scaled asinh-transformed expression of each light chain isotype (colors) for total B cells pooled from three donors before and after peak normalization. **E)** Percent of each B cell subset with a given isotype class label (defined by surface staining of antibody isotypes) for individual donors. ND signifies “not determined” – cells with low/absent expression of all isotypes. IgMD denotes co-expression of IgM and IgD. **F)** Total B cells were segregated by heavy chain isotype. For each isotype, the percent of cells labeled as transitional/naïve, memory/plasma, and ungated was determined for the canonical gating scheme (left) and the meta-clustering classification (right) for individual donors. **G)** Percent of total B cells with a given isotype class label (defined by surface staining of antibody isotypes) for individual donors. ND signifies “not determined” – cells with low/absent expression of all isotypes. IgMD denotes co-expression of IgM and IgD. **H)** Representative contour plot from one donor from the surface screen of total CD45+ cells showing low co-occurrence of IgE and CD19.

**Figure S3:**
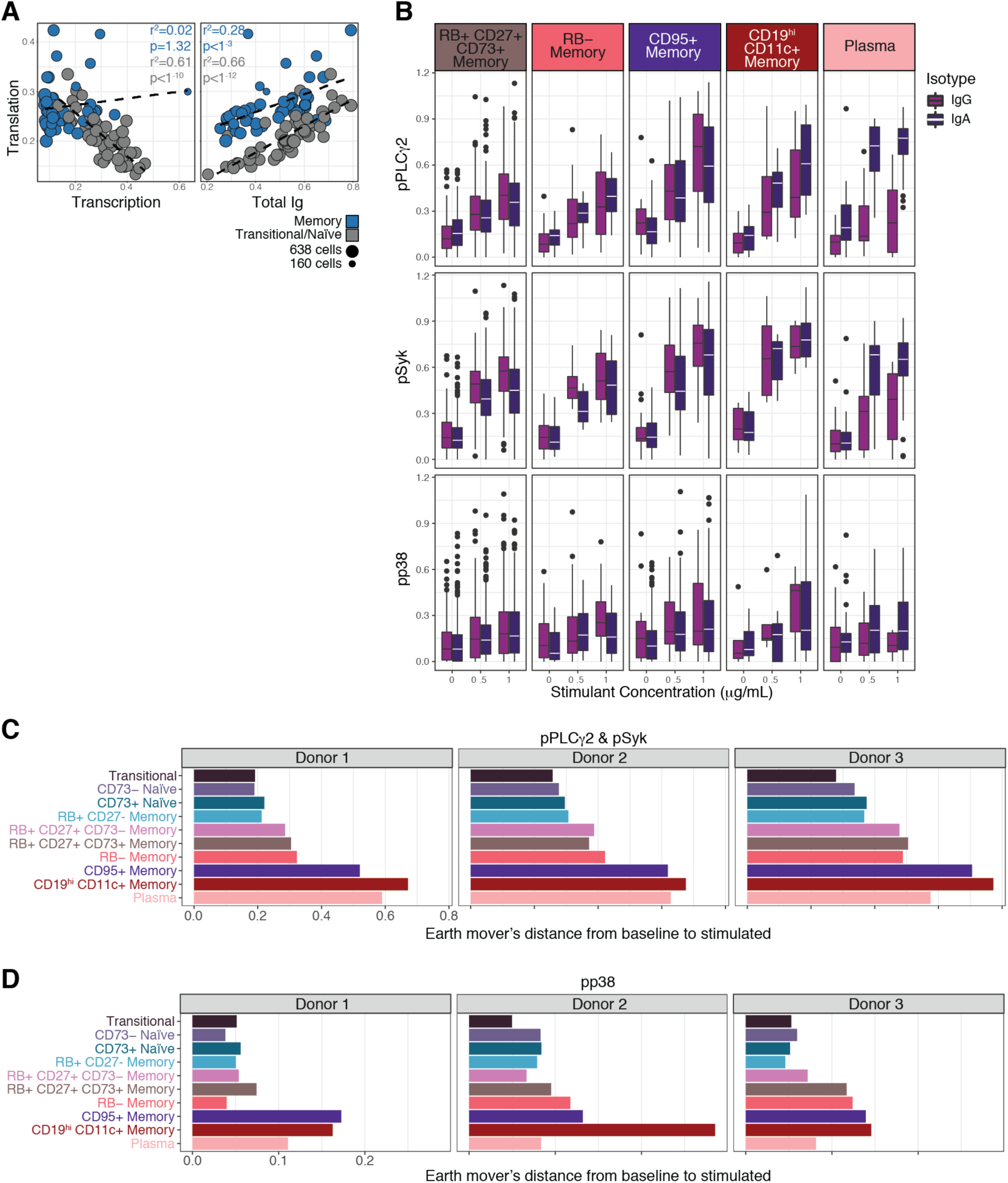
Biosynthesis correlations, mature isotype signaling responses, and individual donor signaling responses – related to Figure 5. **A)** Biaxial plots of median expression of clusters (generated using FlowSOM – also used for initial over-clustering before subset assignment), colored as either memory (blue) or transitional/naïve (grey). Circle size indicates number of cells in cluster. Statistics and lines were calculated from simple linear regression models. **B)** Boxplots of the expression of three signaling molecules at three different doses (x-axis) segregated by isotype (colors) and by phenotype (columns). Only phenotypes that were > 3% IgG+ and > 3% IgA+ were included in visualization. Lines indicate median value, boxes indicate interquartile range (IQR), whiskers indicate first/third quartile +/- 1.5 IQR. Dots indicate outliers. **C)** Quantification of earth mover’s distance from baseline samples to stimulated samples (1 μg/mL) for pPLCγ2 and pSyk for individual donors. **D)** Quantification of earth mover’s distance from baseline samples to stimulated samples (1 μg/mL) for pp38 for individual donors.

